# Preventing cancer by long-term partial Myc suppression

**DOI:** 10.1101/2020.12.30.422650

**Authors:** Nicole M. Sodir, Luca Pellegrinet, Roderik M. Kortlever, Yong-Won Kwon, Shinseog Kim, Daniel Garcia, Alessandra Perfetto, Panayiotis Anastasiou, Lamorna Brown Swigart, Trevor D. Littlewood, Gerard I. Evan

**Affiliations:** Department of Biochemistry, University of Cambridge, 80 Tennis Court Road, Cambridge CB2 1GA, UK; Abcam, 860 Auburn Ct, Fremont, CA 94538, USA; Center for Genomic Integrity, Institute for Basic Science, Ulsan, 44919, Republic of Korea; Oncogenesis Thematic Research Center at Bristol Myers Squibb, 10300 Campus Point Dr, San Diego, CA 92121, USA; Department of Laboratory Medicine, University of California, San Francisco, 2340 Sutter Street, San Francisco, CA 94115, USA

**Keywords:** cancer prophylaxis, Myc, lung cancer, pancreatic cancer

## Abstract

*Myc^+/−^* haploinsufficient mice and mice with deleted *Myc* enhancers express reduced levels of the pleiotropic transcription factor Myc. Such mice are viable, with relatively mild pathologies, but show delay in onset of certain cancers. However, many phenotypes arising from germline gene perturbation are indirect consequences of adaptive developmental compensation and so do not translate to equivalent phenotypes when applied to adults. To ascertain whether systemic Myc hypomorphism also conferred cancer protection when acutely imposed in adults, and what the side-effects of this might be, we constructed a genetically engineered mouse model in which Myc expression may be systemically and reversibly hypomorphed at will. Acute imposition of Myc hypomorphism in adult mice conferred potent protection against both KRas^G12D^-driven lung and pancreatic cancers yet elicited only mild haematopoietic side effects. These side effects were completely suppressed by imposing Myc hypomorphism metronomically – a regimen that nonetheless retained potent cancer prophylaxis. Our data identify a key bottleneck at the transition from pre-malignant hyperplasia to overt tumour that is peculiarly reliant on levels of Myc that are higher than those required for most adult physiology, offering a possible window of opportunity for Myc inhibition in cancer prophylaxis.

## INTRODUCTION

Cancers are extremely heterogeneous diseases that arise through the progressive accumulation in somatic clades of diverse mutations that perturb signalling networks that regulate cell growth, differentiation, repair, survival and movement. In normal cells, such signalling networks preside over tissue ontogeny, maintenance and repair and have consequently evolved to be inherently robust. This robustness is reflected in the multiplicity, diversity and functional redundancy exhibited by the driver mutations that underpin spontaneous cancers, and in the adaptability and evolvability of cancer cells that thwart the pharmacological selective pressures we impose upon them. Nevertheless, substantial evidence now suggests that the integrated output of these robust signalling networks is ultimately funnelled through a small number of non-redundant downstream effectors. One of these is Myc, a highly pleiotropic basic helix–loop–helix leucine zipper transcription factor that serves as a central and obligate conduit to transcriptionally coordinate the diverse intracellular and extracellular programmes necessary for orderly cell and tissue growth and repair ^1–4^.

As a pivotal arbiter of cell proliferation, Myc harbours great oncogenic potential. However, Myc activity in normal cells is tightly restrained by negative feedback transcriptional autoregulation, which limits the peak intracellular level of Myc ^5–7^, and by multiple tiers of auto-attenuation that constrain Myc persistence: both Myc mRNA and protein are very short-lived, expressed only in receipt of mitogenic signalling and, are rapidly cleared from cells upon surcease of mitogenic signals ^8,9^. This tight quantitative and temporal regulation is disrupted by diverse mechanisms in virtually all cancers including amplification of the c-*myc* gene (hereafter *Myc*) or its regulatory enhancers, chromosomal translocation stabilizing mutation and, most commonly, relentless induction by “upstream” oncogenic drivers ^10^. These all result in both accumulation and aberrant persistence of Myc, although it remains unclear which of these is the principal contributor to its oncogenic activity. Myc is frequently overexpressed in many advanced solid malignancies and it has been suggested that such pathologically elevated Myc engages low affinity gene regulatory elements that co-opt novel target genes with oncogenic properties ^11,12^. On the other hand, several recent studies using switchable genetic mouse cancer models indicate that the aberrant persistence of Myc expressed at normal physiological levels and absent over-expression is sufficient to drive all of Myc’s diverse oncogenic activities ^13–15^, which suggests that Myc oncogenic activity is a consequence of pathological persistence of its normal physiological functions. Consistent with this, many studies in diverse genetically engineered mouse cancer models agree that Myc activity is essential for maintenance of diverse cancers, irrespective of their cell type of origin or underlying oncogenic mechanism and, importantly, whether or not Myc is itself a *bona fide* “driver.”

In addition to its acknowledged generic role in maintaining diverse cancers, evidence also suggests that factors governing physiological levels of endogenous Myc play a critical role in cancer predisposition. Normal Myc expression is influenced by a plethora of evolutionarily conserved upstream and downstream enhancer and superenhancer elements, which tailor Myc expression to the specialized developmental, maintenance and regenerative needs of different tissues and organs. However, certain naturally occurring allelic variations within these regulatory regions generate cryptic recognition elements for lineage-specific transcription factors, augmenting Myc expression and greatly predisposing to certain cancers, including colorectal, prostate, bladder and breast cancer and various leukaemias such as ALL and CLL ^16–22^. Furthermore, topological reorganization ^23,24^ and Myc amplification ^25,26^ of *Myc* enhancers are frequent mutagenic events in certain cancer types. Remarkably, wholesale germline deletion of many of these evolutionarily conserved cancer-predisposing regulatory elements markedly decreases incidence of their associated cancers yet has little impact on mouse development or adult tissue physiology ^17,25,27,28^. In many, but not all, instances the observed cancer protection correlates with reduced basal levels of Myc expression in the affected organ ^17^. Myc haploinsufficiency, which similarly reduces basal Myc expression in many tissues ^29,30^, likewise confers substantial protection against onset of intestinal polyposis in *APC^Min^* mice ^31,32^ and KRas^*G12D*^-induced pancreatic adenocarcinoma ^33^. Furthermore, immortalized *Myc* haploinsufficient fibroblasts show marked resistance to Ras or Raf transformation *in vitro* ^34^ as well as reduced global protein and ribosome synthesis, extension of G1 and G2 cell cycle phases ^35^, and an increased tendency to upregulate the CDK4/6 inhibitor p16^*INK4*^ and engage replicative senescence ^36^.

The remarkable suppression of transformation and delay in onset of certain cancers afforded by global reduction of endogenous Myc in enhancer-deleted and haploinsufficient Myc mice suggests the existence of a bottleneck in early tumour evolution whose transit is critically dependent on full physiological levels of endogenous Myc expression. Even though no specific pharmacological inhibitor of Myc has yet been devised, this nonetheless raises the intriguing notion that pharmacological blunting of Myc might have utility in generic cancer prevention. However, caution is needed. First, adult phenotypes arising from germline perturbations of specific genes are seldom solely the direct result of the target gene’s perturbation but often arise from organism-level adaptive developmental compensation to that germline perturbation ^37–39^. Second, Myc is subject to potent homeostatic transcriptional autosuppression ^5–7^. This not only serves to limit peak Myc expression but also governs the temporal dynamics of Myc induction in response to upstream mitogenic signalling, an important consideration for a transcription factor whose activities in normal adult physiology must be tightly confined to the episodic and transient proliferative needs of tissue maintenance and repair. Hence, it is by no means certain that the cancer protection conferred by germline Myc hypomorphism would translate into equivalent protection afforded by acutely-induced Myc reduction in adults. Anyway, Myc germline haploinsufficient mice present with unwelcome side-effects, including small body size with variably reduced cellularity across organs ^30^, infertility ^29^, and compromised haematopoiesis due to aberrant self-renewal of haematopoietic stem cells ^40^. The extent to which these pathologies are direct consequences of Myc hypomorphism versus results of developmental compensation is, again, unknown. Therefore, to model acute pharmacologically-imposed Myc hypomorphism in adults we constructed a novel mouse in which hypomorphism of endogenous Myc may be reversibly induced and relieved at will. We used this *in vivo* model to assess the cancer prophylactic efficacy of acute Myc hypomorphism and to determine any iatrogenic adverse impacts that such partial systemic Myc inhibition might elicit.

## RESULTS

To determine whether imposition of endogenous Myc hypomorphism in adults also offers protection against cancers, we generated a mouse in which endogenous Myc expression may be reversibly hypomorphed at will. We adapted technology previously developed for ectopic regulation of the endogenous *e2f3* gene ^41^ in which a heptameric tetracycline-response element (*TRE*) is inserted into the endogenous gene target. This *TRE* then serves as a recognition motif for the tetracycline-dependent tTS^*Kid*^ transcriptional repressor ^42,43^ rendering the endogenous target gene susceptible to rapid and complete repression upon removal of tetracycline. We reasoned that targeting the *TRE* away from the endogenous *Myc* promoter and instead to the downstream 2^nd^ intron (Supplementary Figure 1A) would allow for ectopically regulatable partial repression of endogenous *Myc*. The resulting *Myc^TRE/TRE^* (*M*) mice were crossed with our *tTS^Kid^* strain (*R*), in which the tTS^*Kid*^ repressor is ubiquitously and constitutively expressed from the β-actin promoter ^41^, to generate *Myc^TRE/TRE^*; *tTS^Kid/−^* (hereafter called *MR*) mice. In the absence of tetracycline, we hypothesized the tTS^*Kid*^ repressor should bind to the 2^nd^ *intron TRE* and partially repress *Myc* expression. However, administration of tetracycline should rapidly and reversibly relieve this partial repression – in effect generating a reversibly switchable Myc hypomorph (Supplementary Figure 1B).

To validate reversible Myc hypomorphism in *MR* mice *in vivo*, we examined levels of endogenous Myc expression in adult tissues that normally proliferate and so have measurable constitutive Myc expression - bone marrow, spleen, thymus and small intestine. Wild type levels of Myc were maintained in *MR* mice throughout gestation and into adult life by sustained administration of tetracycline. Tetracycline was then withdrawn to activate the *tTS^Kid^* repressor and Myc RNA levels assessed by RT-PCR over the following 4 weeks. Representative data for small intestine, spleen, thymus and bone marrow show a profound, stable and reproducible partial decline in endogenous *Myc* expression of between 20 and 50%, depending on tissue, which is completely reversed within 1 week upon re-addition of tetracycline (Supplementary Figure 2A). During the 4-week test period of systemic Myc hypomorphism we observed no detectable deleterious impact on animal health, activity or welfare, or tissue architecture (Supplementary Figure 2B) nor any significant perturbation in blood cell counts (Supplementary Figure 2C).

Determining the extent of Myc hypomorphism in most other *MR* cancer-sensitive adult tissues, for example lung and pancreas, is complicated by the fact that they are usually non-proliferative and consequently express negligible levels of endogenous Myc. Nonetheless, both lung and pancreas undergo extensive proliferation and growth during early postnatal life along with concomitant expression of endogenous Myc. Therefore, we assessed the efficacy of switchable Myc hypomorphism in neonatal *MR* lung and pancreas. *MR* embryos were allowed to develop with normal Myc levels and tetracycline withdrawn at birth to hypomorph Myc. 8 and 14 days later, hypomorphed neonates were euthanized, lungs and pancreata harvested, and endogenous *Myc* mRNA levels assessed (Supplementary Figure 3A). Relative to controls (animals maintained on tetracycline), lung and pancreas of hypomorphed neonatal mice showed profound partial Myc mRNA repression (from 30-60%) at both time points (Supplementary Figure 3B).

Since germline Myc hypomorphism significantly delays onset of tumours in various cancer mouse models, we next asked whether Myc hypomorphism acutely imposed on adult mice is similarly protective. We crossed *MR* mice into the well-characterized *LSL-Kras^G12D^* mouse model of lung adenocarcinoma (LUAD) ^44^ to generate adult *MRK* mice. Myc was then hypomorphed (tetracycline-deprived) for 3 weeks in adult mice and KRas^*G12D*^ then sporadically activated in lung epithelium by Adenovirus-Cre inhalation. Thereafter, the mice were maintained in a hypomorphic state, then euthanized 12, 22 or 32 weeks later and tumour loads assessed (Figure 1A). As expected, control, non-hypomorphed mice maintained on tetracycline exhibited a progressively worsening tumour burden over 32 weeks, together with increasing representation of high grade, aggressive disease, including adenocarcinomas. By contrast, lungs of hypomorphed animals exhibited multiple early stage hyperplastic (AAH) lesions but progression to frank tumour was completely absent in the hypomorphed animals aside from a very few grade-1 lesions (Figure 1B). Thus, adult imposed Myc hypomorphism has no discernible inhibitory impact on the genesis of pre-malignant KRas^*G12D*^-induced lung AAH but potently blocks the transition and growth of these pre-malignant lesions into overt malignant lung tumours.

**Figure 1:**
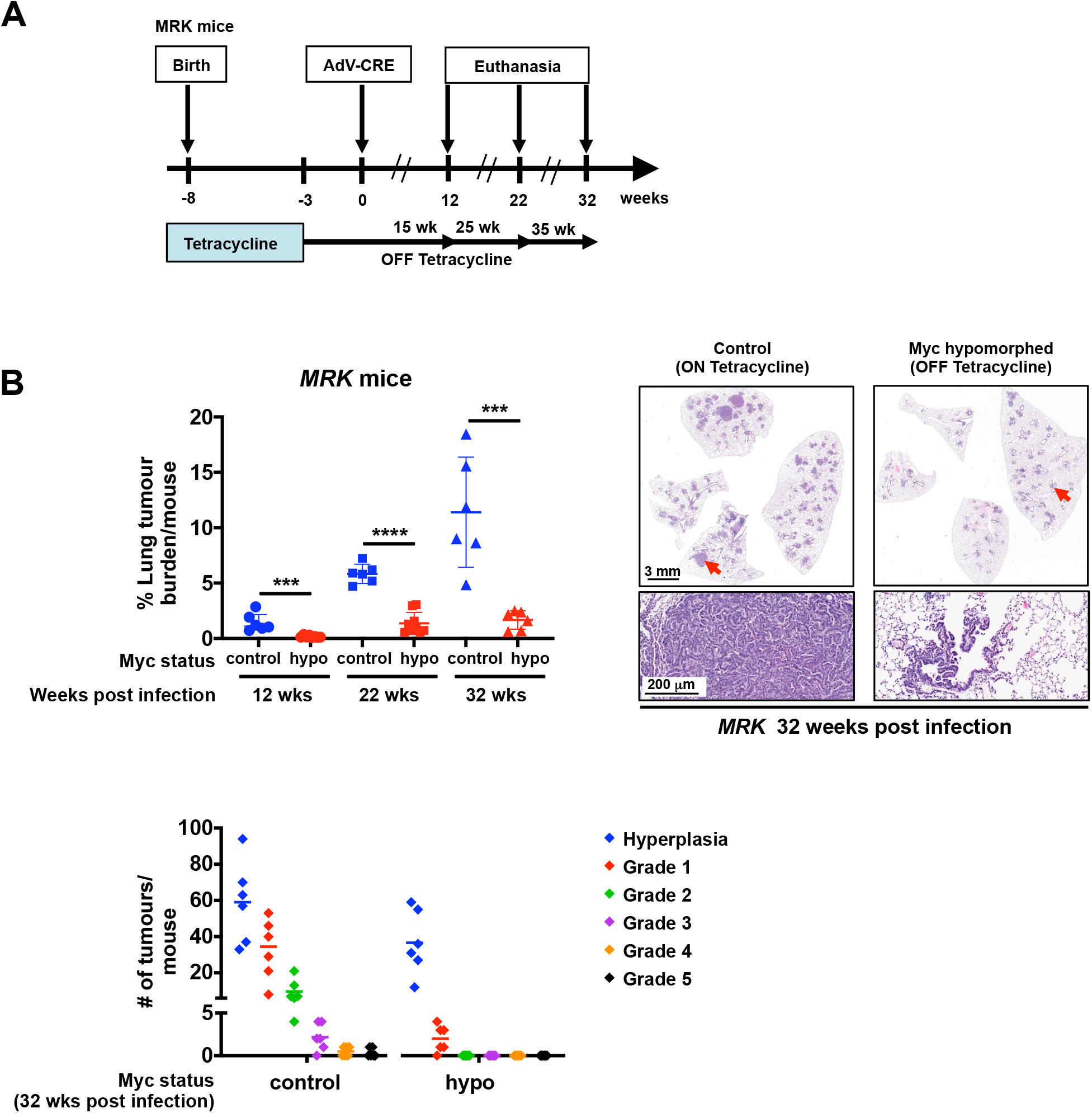
Imposition of Myc hypomorphism in adult mice blocks progression of KRas^*G12D*^-driven lung hyperplasia to adenocarcinoma *in vivo*. **A.** Schematic of study. Endogenous Myc was maintained at normal levels in *MRK* (*Myc^TRE/TRE^;β-actin-tTS^Kid/−^;LSL-KRas^G12D/+^*) mice through development, neonatal growth and into adulthood (5 weeks old) by continuous administration of tetracycline. Endogenous Myc was then hypomorphed by withdrawal of tetracycline and Adv-Cre administered by inhalation three weeks later to trigger sporadic expression of KRas^*G12D*^ in lung epithelium. Mice were then euthanized 12, 22 or 32 weeks later. Control mice were treated identically save that tetracycline was maintained throughout. **B.** Top left: Percentage of tumour burden relative to total lung in non-hypomorphed versus hypomorphed *MRK* mice at 12, 22 or 32 weeks post Adv-Cre activation of *KRas^G12D^*. Results depict mean ± SD in each treatment group. The unpaired t-test was used to analyze tumor burden. ***p < 0.001, ****p < 0.0001. SD = standard deviation. Top right: Representative H&E-stained sections of lungs from *MRK* mice at 32 weeks post Adv-Cre activation of *KRas^G12D^* either non-hypomorphed (maintained on tetracycline) or hypomorphed (off tetracycline). Arrows mark regions shown at higher magnification below. Bottom left: Grading of tumours (following ^44^) in lungs from *MRK* mice at 32 weeks post Adv-Cre activation of *KRas^G12D^* either non-hypomorphed (maintained on tetracycline) or hypomorphed (off tetracycline). Results depict quantitation of total numbers of tumours of each grade, with the means indicated.

Inactivation of p53 greatly accelerates KRas^*G12D*^-driven LUAD progression ^45^. We therefore next asked whether p53 inactivation negates the prophylactic impact of Myc hypomorphism on lung tumourigenesis. *MRK* mice were crossed into a *p53^flox/flox^* background to generate *MRKP^fl^* mice. This allowed for concurrent activation of KRas^*G12D*^ and deletion of p53 in lung epithelia upon Adenovirus-Cre inhalation. The previous hypomorph tumour prophylaxis experiment was then repeated. As reported ^45^, p53 inactivation greatly accelerated lung tumour progression in the non-hypomorphed control *MRKP^fl^* animals, requiring them to be culled after only 14 weeks. Nonetheless, Myc hypomorphism once again potently retarded tumour formation and markedly reduced tumour load (Figure 2A). However, in addition to an abundant background of AAH lesions, similar to that we had observed in hypomorphed p53*^wt^ MRK* animals, we also noted the presence of a few sporadic large, invasive lung tumours. Investigation revealed that almost all of these were “escapee” tumours that had bypassed the hypomorphism switch either by losing tTS^*Kid*^ repressor expression or by gross up-regulation of endogenous Myc, or both (Figure 2B). Hence, p53 inactivation does not itself negate the cancer protection afforded by hypomorphing Myc although it does increase the probability of happenstance accidents that break our experimental hypomorph model.

**Figure 2:**
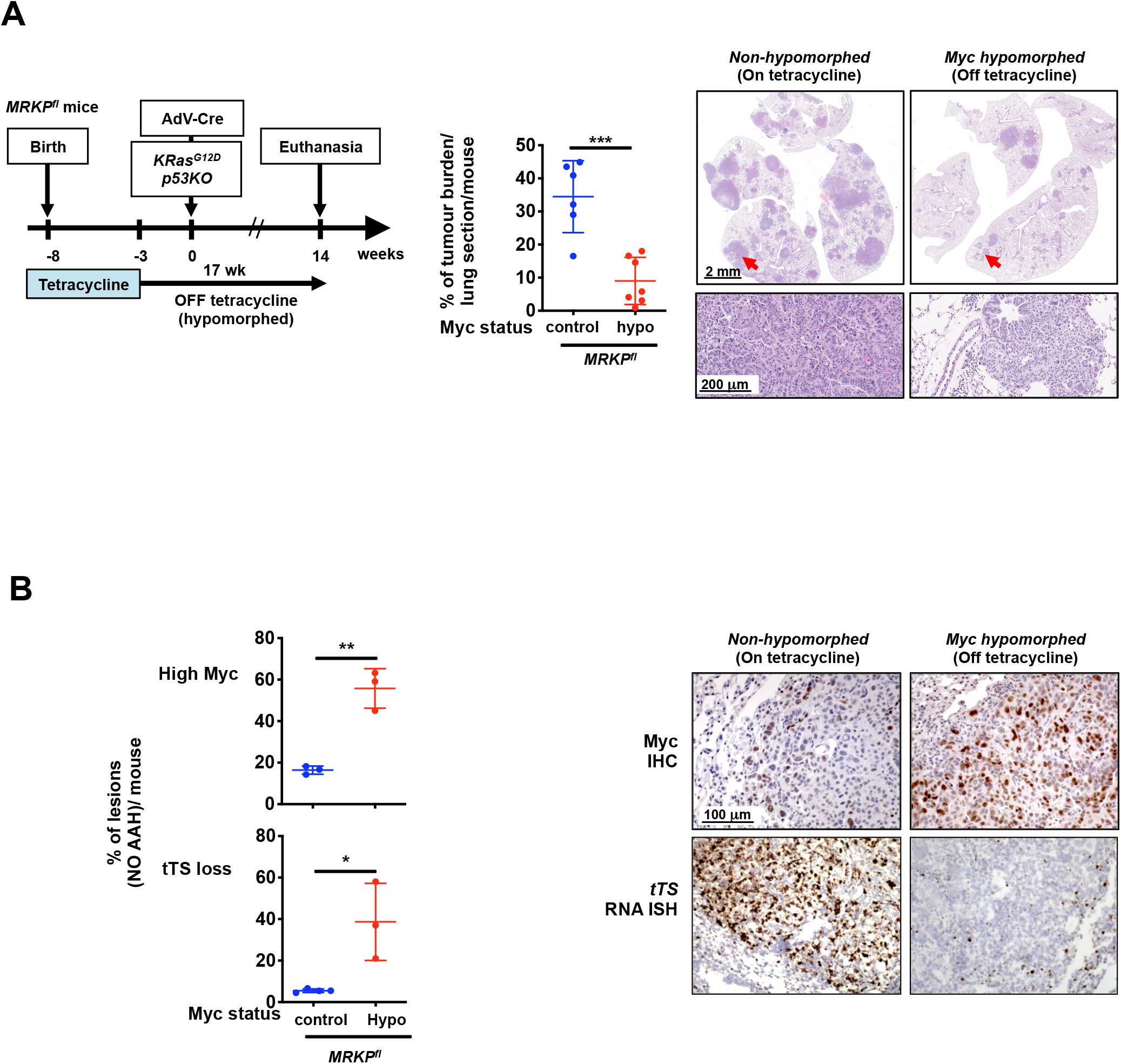
p53 is not required for lung tumour prevention conferred by Myc hypomorphism. However, p53 inactivation does facilitate sporadic mechanisms of tumour escape. **A.** Left: Schematic of study. *MRKP^fl^* (*Myc^TRE/TRE^;β-actin-tTS^Kid/−^;LSL-KRas^G12D/+^;p53^flox/flox^*) mice were maintained on tetracycline (endogenous Myc at *wt* levels) throughout embryonic development until 5 weeks of age. Tetracycline was then withdrawn to hypomorph Myc and 3 weeks later Adv-Cre administered by inhalation to sporadically and concurrently activate expression of KRas^*G12D*^ and inactivate p53 in lung epithelium. Mice were euthanized 14 weeks post Adv-Cre inhalation and lung tissue harvested. Control animals were treated identically save that they were maintained throughout on tetracycline to sustain *wt* endogenous Myc levels. Centre: quantitation of overall tumour load in non-Myc hypomorphed (blue) versus hypomorphed (red) *MRKP^fl^* mice at 14 weeks post Adv-Cre inhalation. Results depict mean ± SD in each treatment group. Right: representative H&E staining of lung tissue harvested from non-hypomorphed versus hypomorphed *MRKP^fl^* mice 14 weeks post Adv-Cre inhalation. Arrows mark regions shown at higher magnification below. **B.** Analysis of mechanisms by which lung tumours in *MRKP^fl^* mice escape Myc hypomorphism. Left: quantification of individual lung tumours exhibiting demonstrably elevated levels of Myc (Myc IHC positive cells >25%) or complete loss of the *tTS^Kid^* repressor expression (RNA ISH) in control non-hypomorphed (Blue) (ON tetracycline) versus Myc hypomorphed (Red) (OFF tetracycline) *MRKP^fl^* mice 14 weeks post Adv-Cre infection. Results represent percentage of total lesions, mean ± SD. Right: top panels show representative images of Myc IHC staining in non-hypomorphed lung tumour (left) versus an example of dramatic Myc upregulation in a large escapee lung tumour from a “hypomorphed” (off tetracycline) *MRKP^fl^* mouse. Bottom panels show RNAscope staining for *tTS^Kid^* repressor expression, which is broadly expressed by lung tumour cells in non-hypomorphed *MRKP^fl^* mice (left) but absent from an escapee lung tumour in a *MRKP^fl^* mouse in the absence of tetracycline (right). The unpaired t-test was used to analyze data. *p < 0.05, **p < 0.01, ***p < 0.001. SD = standard deviation.

We next asked whether Myc hypomorphism similarly protects against aggressive pancreatic cancers using the well-characterized *LSL-KRas^G12D/+^*;*LSL-p53^R172H/+^*;*pdx1-Cre;* (*KPC*) mouse model of pancreatic adenocarcinoma (PDAC) ^46^. In this model, Cre recombinase expression, driven from the *pdx/IPF1* (*Pancreas/duodenum homeobox protein 1*) promoter in embryonic pancreatic and duodenal progenitor cells from around E8, concurrently triggers expression of both KRas^*G12D*^ and the gain-of-function mutant p53^R172H 46^. *KPC* mice rapidly develop invasive and metastatic PDAC that closely recapitulates all characteristics of the human disease. *KPC* mice were crossed into the *MR* background to generate *MRKPC* mice and tetracycline then withdrawn at 4 weeks of age to hypomorph Myc expression. Tumours were allowed to develop and after 18 weeks mice were euthanized and pancreata isolated and assessed for tumour load. Pancreata of most (~85%) control (non-hypomorphed) mice exhibited an extremely high tumour load comprising multiple invasive pancreatic adenocarcinomas. By contrast, the pancreata of hypomorphed Myc-mice showed dramatically reduced tumour burden formed almost exclusively of indolent PanINs (Figure 3A). However, just as with the lungs of *MRKP^fl^* mice (above), we noted occasional large aggressive pancreatic tumours in some 30% of Myc-hypomorphed animals. Once again, analysis of these “escapees” showed they had markedly upregulated expression of endogenous Myc and lost tTS^*Kid*^ repressor expression (Figure 3B) – happenstance mechanisms that break the mechanism of our switchable hypomorphism Myc model.

**Figure 3:**
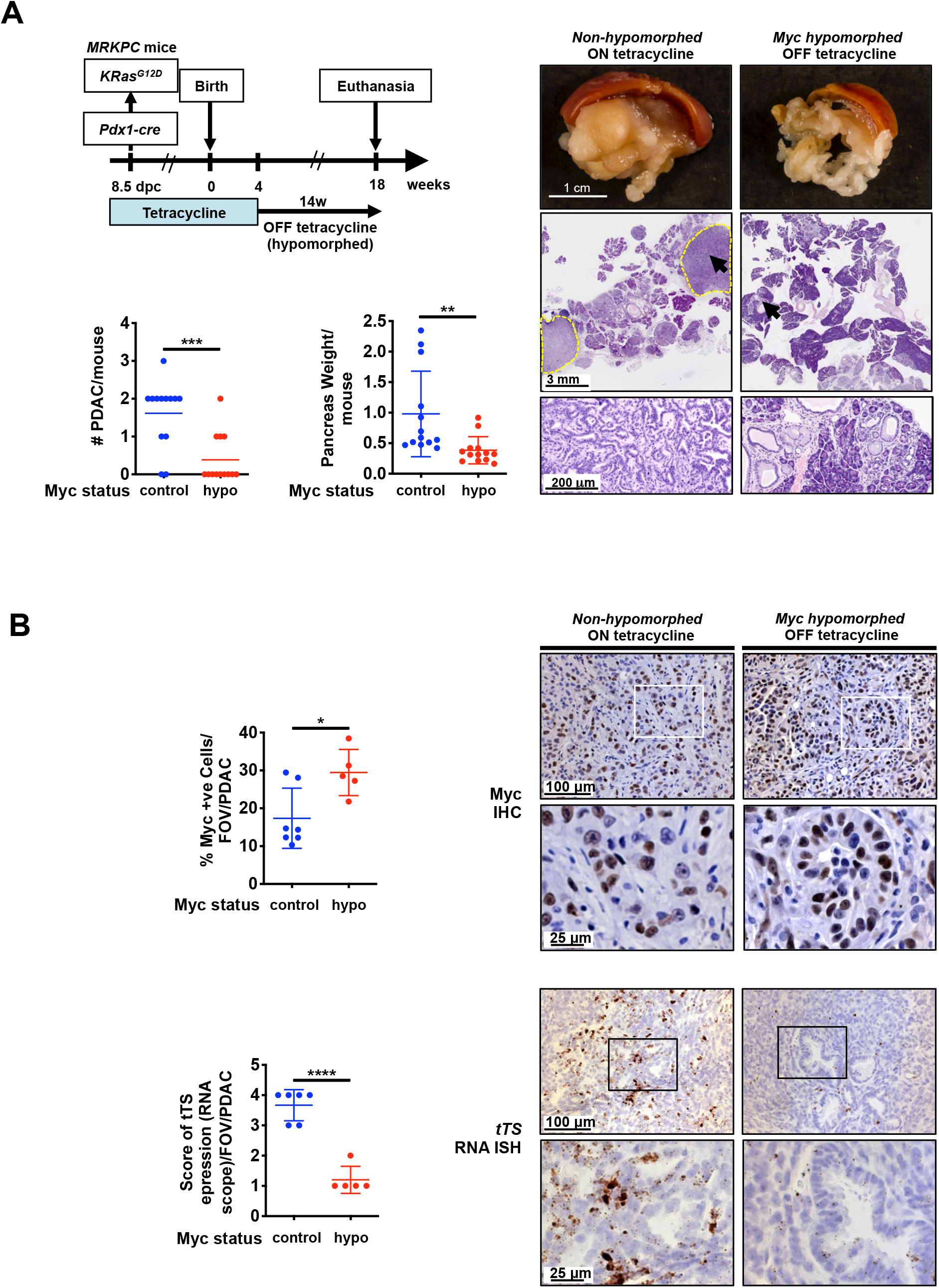
p53 is not required for pancreas tumour prevention conferred by Myc hypomorphism. However, p53 inactivation does facilitate sporadic mechanisms of tumour escape. **A.** Upper left: Schematic of study. In *MRKPC* (*Myc^TRE/TRE^;β-actin-tTS^Kid/−^;LSL-KRas^G12D/+^;LSL-p53^R172H/+^;Pdx-1-Cre*) mice expression of Cre recombinase is driven from the *pdx/IPF1* promoter, triggering co-expression of KRas^*G12D*^ and p53^*R172H*^ in pancreatic progenitor cells from around 8.5 *dpc*. *wt* levels of endogenous Myc were maintained throughout development and into adulthood by continuous administration of tetracycline. At 4 weeks of age, tetracycline was withdrawn from mice to hypomorph endogenous Myc and animals euthanized 14 weeks later (total 18 weeks old). Tetracycline administration was maintained throughout in the control, non-hypomorphed cohort. Bottom left: quantitation of PDAC tumours per mouse in non-hypomorphed (blue) versus hypomorphed (red) *MRKPC* mice and overall pancreas weight (a surrogate for tumour load) in the same animals. Right: representative macroscopic, low- and high-power images of non-hypomorphed versus hypomorphed pancreata at 18 weeks, showing multiple PDAC tumours in the former and none in the latter. Dotted yellow lines represent the margins of a PDAC. Arrows mark regions shown at higher magnification below. **B.** Analysis of mechanisms by which tumours escape in Myc hypomorphed *MRKPC* mice. Top left: quantitation of cells positive for Myc expression (IHC) in PDACs from non-hypomorphed *MRKPC* mice (blue) versus escapee PDACs from hypomorphed counterparts (red). Top right: Myc IHC staining in representative examples of PDACs from non-hypomorphed *MRKPC* mice (left) versus hypomorphed PDAC escapee (right). Boxes represent regions shown at higher magnification below. Bottom left: quantitation of RNAscope analysis of *tTS^Kid^* expression in PDACs from non-hypomorphed *MRKPC* mice (blue) versus a “hypomorphed” PDAC escapee (red). The *tTS^Kid^* score for each PDAC was calculated as follows: score 1 = <25% of PDAC cells express *tTS^Kid^* RNA, score 2 = 25% = 50% of PDAC cells express *tTS^Kid^*, score 3 = >50% to 75% of PDAC cells express *tTS^Kid^*, score 4 = >75% of PDAC cells express *tTS^Kid^*. Bottom right: representative RNAscope *in situ* hybridization analysis showing *tTS^Kid^* expression in PDAC from non-hypomorphed (no tetracycline) *MRKPC* mice and its absence from an escapee PDAC tumour (+tetracycline). Boxes represent regions shown at higher magnification below. The unpaired t-test was used to analyze the data. Mean ± SD are shown. *p < 0.05, **p < 0.01, ***p < 0.001, ****p < 0.0001. FOV = field of view. SD = standard deviation.

Our data demonstrate that acute adult imposition of relatively mild repression of endogenous Myc acts as a powerful block to the evolution of overt cancers. To map the precise location of the Myc level-dependent bottleneck in progression of the lung and pancreas cancers we constructed an allelic series of knock-in mice in which different levels of the reversibly switchable, 4-hydroxytamoxifen-dependent Myc variant MycER^T2^ may be activated at will in lung or pancreas epithelium. We previously showed that MycER^T2^ expression driven by two alleles of the endogenous *Rosa26* promoter (*R26^MT2/MT2^* homozygotes) is at a level broadly equivalent to that of endogenous Myc in proliferating cells and tissues ^13,15^. By contrast, mice in which MycER^T2^ expression is driven from only a single *Rosa26* allele (*R26^MT2/+^* hemizygous) express only half as much MycER^T2 13^, a sub-physiological level broadly similar to that of hypomorphed Myc in proliferating *MR* tissues. As previously shown, acute activation of physiological levels of MycER^T2^ in (*LSL-KRas^G12D/+^*; *R26^MT2/ MT2^*-referred to as *KR26^MT2/MT2^*) lung and (*LSL-KRas^G12D/+^*;*pdx1-Cre; R26^MT2/MT2^*-referred to as *KCR26^MT2/MT2^*) pancreas *in vivo* is sufficient to drive full oncogenic cooperation with oncogenic KRas^*G12D*^, triggering immediate and synchronous transition of indolent KRas^*G12D*^-driven lung pre-tumours (Figure 4A) and PanINs (Figure 4B) to adenocarcinomas with all the signature invasive, proliferative, stromal, inflammatory, immune and vascular attributes of spontaneous adenocarcinomas from those same organs ^14,15^. By contrast, activation of hypomorphic levels of MycER^T2^ in the indolent tumours of hemizygous (*LSL-KRas^G12D/+^*; *R26^MT2/+^*, referred to as K*R26^MT2/+^*) lung and (*LSL-KRas^G12D/+^*; *pdx1-Cre; R26^MT2/+^*, referred to as *KCR26^MT2/+^*) elicited no measurable change in lung adenoma/PanIN morphology, proliferation or stroma (Figure 4A&B). Taken together, our data demonstrate that modest Myc hypomorphism, imposed in adult animals, powerfully suppresses KRas^*G12D*^-driven lung and pancreas cancer, most probably by blocking the key transition from indolent to malignant neoplasia.

**Figure 4:**
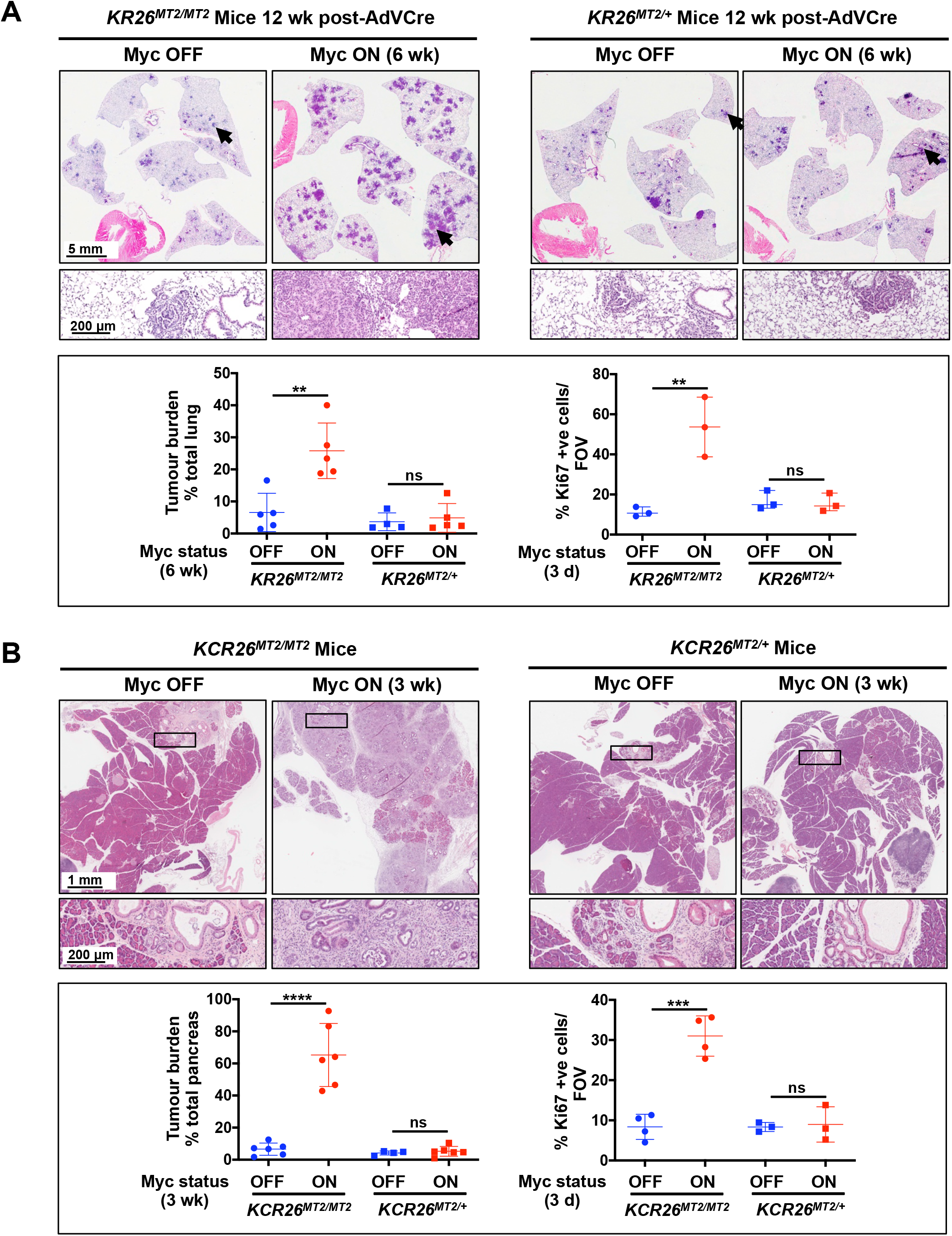
A tight minimum threshold of Myc activity is required to drive transition from indolent pre-tumour to adenocarcinoma *in vivo*. **A.** Adenovirus-Cre was administered by inhalation to either *KR26^MT2/MT2^* (*LSL-KRas^G12D/+^*;*R26^MT2/MT2^*) or *KR26^MT2/+^* (*LSL-KRas^G12D/+^*;*R26^MT2/+^*) adult mice to trigger sporadic expression of KRas^*G12D*^ and MycER^T2^ in lung epithelia. 6 weeks later (i.e. after 6 weeks of KRas^*G12D*^-only activity) MycER^T2^ was also activated for a further 6 weeks (Myc ON). MycER^T2^ was not activated in control mice (Myc OFF). Representative H&E-stained sections of lungs from *KR26^MT2/MT2^* versus *KR26^MT2/+^* mice are shown without (left) and with (right) Myc activation. H&E panel inserts below show arrowed regions at higher magnification. Lower panel left: quantitation of percentage tumour burden relative to the total lung in *KR26^MT2/MT2^* and *KR26^MT2/+^* mice, either without or with Myc for 6 weeks. Results depict mean ± SD, n = 5 mice per treatment group. Lower panel right: quantitation of proliferation (immunohistochemical staining for Ki67) in sections of lungs harvested from *KR26^MT2/MT2^* and *KR26^MT2/+^* mice 12 weeks after activation of KRas^*G12D*^ either without or with activation of MycER^T2^ for 3 days. Results depict mean ± SD, n = 3 mice per treatment group **B.** Representative H&E-stained sections of pancreata from 15 week-old *KCR26^MT2/MT2^* (*LSL-KRas^G12D/+^*; *pdx1-Cre;R26^MT2/MT2^*) and *KCR26^MT2/+^* (*LSL-KRas^G12D/+^*;*pdx1-Cre; R26^MT2/+^*) mice either without or with Myc activation for 3 weeks. H&E panel inserts below show boxed regions at higher magnification. Lower panel left: quantitation of percentage tumour burden relative to the total pancreas from *KCR26^MT2/MT2^* and *KCR26^MT2/+^* mice, either without or with Myc activation for 3 weeks. Results depict mean ± SD, n = 4-6 mice per treatment group. Lower panel right: quantitation of proliferation (Ki67) in sections of pancreata harvested from 12 week-old *KCR26^MT2/MT2^* and *KCR26^MT2/+^* mice, either without or with activation of MycER^T2^ for 3 days. Results depict mean ± SD, n = 3-4 mice per treatment group. Data were analyzed using unpaired t-test. **p < 0.01, ***p < 0.001, ****p < 0.0001, ns= non-significant. SD = standard deviation. Part of figure 4 (some data for *KR26^MT2/MT2^* and *KCR26^MT2/MT2^* mice) is included for reference from previous publications^14,15^.

Myc is a transcription factor that exerts its oncogenic impacts through its regulation of target genes. To address whether the reduced MycER^T2^ levels in heterozygous *R26^MT2/+^* tissues can still generate transcriptional output, we assessed induction of the Myc target gene *Rasd2* (TRANFAC Curated Transcription Factor Targets Dataset) in pancreata of 12 week-old homozygous *KCR26^MT2/MT2^* versus hemizygous *KCR26^MT2/+^* mice. Although reduced relative to the levels induced in homozygous *R26^MT2/MT2^* tissue, induction of *Rasd2* was nonetheless clearly detectable in heterozygous *R26^MT2/+^* pancreas (Supplementary Figure 4).

A major concern with any future exploitation of long-term Myc suppression in cancer prophylaxis is that Myc is required for many process during development, neonatal growth, and adult maintenance and repair of many adult tissues including lung ^47^, pancreas ^48–51^ and bone marrow ^52–54^. Homozygous germline *Myc* knockout embryos fail around 10.5 *dpc* ^29,30,55^, which coincides temporally with onset of extensive embryonic vasculogenesis and haematopoiesis ^56^. And while haploinsufficent Myc hypomorphic mice survive development and are born at normal Mendelian frequency, they grow to significantly smaller adult size with variable levels of reduced size and cellularity across their tissues ^30,57^ and a variety of pathologies, as already discussed.

To search for any pathologies associated with acutely imposed hypomorphism in *MR* mice we first asked whether it is compatible with embryonic development. *Myc^TRE/TRE^* (*M*) females were mated with *MR* (*Myc^TRE/TRE^*;*tTS^Kid/−^*) males and maintained either with or without tetracycline. *MR* embryos carried by *M* mothers maintained on tetracycline (i.e. *wt* Myc levels) were born at a normal Mendelian ratio and developed normally, exactly like their *M* (*Myc^TRE/TRE^;tTS^Kid^* negative) control littermates. By contrast, *MR* embryos deprived of tetracycline (i.e. Myc hypomorphed) all failed and were resorbed by 13.5 *dpc* (Supplementary Figure 5A and B). Embryo failure appeared due to a range of developmental delays, most obviously in vasculogenesis. Unexpectedly, this mid-gestation lethality was only evident when the breeding male, not the female, contributed the *tTS^Kid^* repressor (i.e. **♀***M* x ♂*MR*). In crosses where the *tTS^Kid^* repressor allele was provided by the mother (i.e. **♀***MR* x ♂*M*), Myc hypomorphism triggered immediate failure and resorption of both hypomorphed and *wt* embryos. Since this happened to embryos both with and without the *tTS^Kid^* repressor gene, such early failure must be a consequence of Myc hypomorphism in the mother, not the embryos. We guess such infertility reflects a requirement for maximal Myc expression in early pregnancy and is presumably due to dysfunctional decidualization or endometrial-blastocyst signalling in hypomorphed mothers ^58^. This contraceptive phenotype was rapidly reversed upon re-administration of tetracycline to breeding *MR* females and M males, generating *MR* pups at their expected Mendelian ratios (Supplementary Figure 5C).

We next investigated the impact of imposing Myc hypomorphism from birth on postnatal growth and organ maturation. *MR* embryos were allowed to develop with normal Myc levels and tetracycline then withdrawn at birth to hypomorph Myc. 8 and 14 days later, neonates were euthanized, lungs and pancreata harvested, and tissues examined histologically and for proliferation (Ki67) relative to non-hypomorphed controls. Despite substantial reductions in endogenous Myc expression at both 8 and 14 days post-partum (Supplementary Figure 3B), both organs showed identical, normal architecture and proliferation indices (Supplementary Figure 6A). Furthermore, rates of increase in total animal weight were identical in both hypomorphed and control cohorts, with both hypomorphed and control animals achieving the same final adult size at the same time (Supplementary Figure 6B and data not shown).

To identify temporally the developmental window at which Myc hypomorphism impacts mouse development, *MR* embryos were hypomorphed mid-gestation (starting) by withdrawal of tetracycline from pregnant mothers at 13.5 *dpc* and Myc hypomorphism maintained thereafter. Remarkably, such post 13.5 *dpc* hypomorphism had no measurable deleterious or retarding impact on embryo development, neonatal growth or proliferation, nor any adverse impact on final adult organ size and histology (Supplementary Figure 7). Hence, the 13.5 *dpc* mid-gestation bottleneck that is absolutely dependent on maximal Myc expression is transient.

Any future cancer prophylaxis based on partial Myc inhibition will likely require long-term Myc suppression. We therefore next investigated the impact on adult *MR* mouse homeostasis and health over extended periods of sustained Myc hypomorphism. Starting around 6 weeks, we observed a reproducible and progressive leukopenia, a somewhat later onset of erythropenia, and later still thrombocythemia from around 18 weeks (Supplementary Figure 8A). We also noted modest splenic extramedullary haematopoiesis in ~50% of animals after 15 weeks (Supplementary Figure 8B). While all affected mice remained active and appeared well throughout, this nonetheless demonstrated that maximal expression of endogenous Myc is required for long-term maintenance of optimal adult haematopoiesis. We also noted, however, that relaxation of Myc hypomorphism upon re-administration of tetracycline rapidly restored normal blood counts in all animals (Supp. Fig. 2A and data not shown). We therefore asked whether the adverse long term side effects of sustained Myc hypomorphism could be mitigated by metronomic imposition while still affording cancer protection using the KRas^*G12D*^-driven model of lung adenocarcinoma described earlier (Figure 1). KRas^*G12D*^ was activated in *MRK* mice lung epithelium by inhalation of Adenovirus-Cre recombinase ^44^ and animals thereafter maintained on a regimen of alternating 4-week periods of Myc hypomorphism followed 1 week’s reversal, for a total of 22 weeks (Figure 5A). Metronomically hypomorphed mice blood counts remained normal throughout (Figure 5B) while all pre-neoplastic lesions stayed stalled and tumour load remained negligible, in stark contrast to the fulminant tumours in lungs of control, non-hypomorphed, *MRK* mouse controls (Figure 5C).

**Figure 5:**
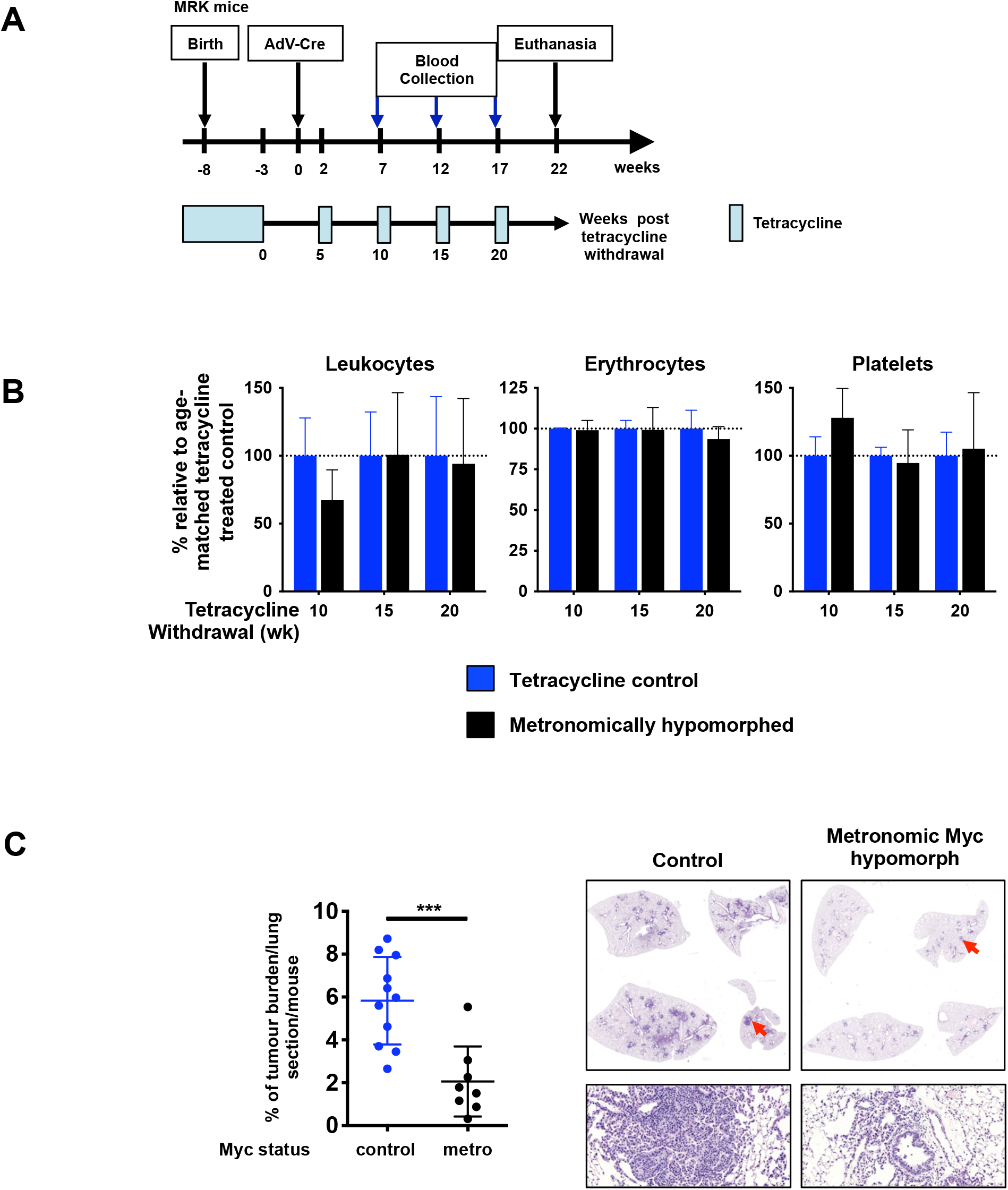
Metronomic Myc hypomorphism circumvents associated haematological pathologies but still protects against KRas^*G12D*^-driven tumourigenesis. **A.** Schematic of metronomic prevention study. Endogenous Myc was maintained in *MRK* mice at normal physiological *wt* levels throughout development and into adult life by continuous administration of tetracycline. At 5 weeks of age tetracycline was withdrawn to hypomorph endogenous Myc and 3 weeks later Adv-Cre administered by inhalation to sporadically induce expression of KRas^*G12D*^ in lung epithelium. Two weeks post Adv-Cre inhalation on, mice were subjected to a metronomic regimen of 1 week’s relaxation followed by 4 weeks’ hypomorphic Myc for a total of 4 cycles. Peripheral blood was withdrawn for analysis just prior to each 1 week’s relaxation time point. Normal levels of endogenous Myc were maintained in control mice by continuous administration of tetracycline throughout. Mice were euthanized 22 weeks post Adv-Cre activation of *KRas^G12D^* and tissues harvested. **B.** Metronomic imposition of Myc hypomorphism spares mice from haematological pathology. Tetracycline was continuously administered to *MRK* mice through development and to 5 weeks of age to maintain physiological Myc expression. Tetracycline was then periodically withdrawn and peripheral blood sampled at the times indicated. Control mice were maintained on tetracycline throughout. Results represent mean ± SD in each treatment group. n≥4 mice per time point. Unpaired t-test shows non-significant difference between groups). **C.** Left: percentage tumour burden relative to total lung in control (normal Myc level – blue) versus metronomically Myc hypomorphed *MRK* mice (black) 22 weeks post Adv-Cre activation of *KRas^G12D^*. The unpaired t-test was used to analyze tumour burden. ***p < 0.001. SD = standard deviation. Right: Representative H&E-stained sections of lungs from control (normal Myc level) versus metronomically Myc hypomorphed *MRK* mice 22 weeks post Adv-Cre activation of *KRas^G12D^*. Arrows mark regions shown at higher magnification below.

## DISCUSSION

In addition to its well documented role in the maintenance of diverse cancer types, circumstantial evidence has mounted that Myc plays a pivotal gatekeeper role in early cancer evolution, irrespective of oncogenic mechanism, cell of origin, or even whether or not Myc is a canonical “driver” mutation. Thus, relatively subtle germline perturbations in *Myc* gene enhancer regulatory elements that govern basal levels of endogenous Myc expression exert huge impacts on lifetime risk of diverse cancers. Most notably, germline modifications such as enhancer deletion and *Myc^+/−^* haploinsufficiency, which reduce basal Myc levels, confer significant resistance to a wide variety of adult-onset cancers while exerting little obvious impact on normal mouse development and adult tissue homeostasis ^17,27^. Such observations raise the intriguing notion that active partial suppression of Myc in adults might likewise serve as a general cancer prophylactic. However, many germline phenotypes fail to translate into corresponding therapeutic outcomes: some are locked in during embryogenesis or neonatal tissue maturation while others are indirect consequences of systemic compensation. Consequently, the extent to which acute imposition of Myc hypomorphism in adults might have a similar prophylactic impact while enjoying a similar paucity of side effects is uncertain.

To address these issues, we developed a novel mouse model in which endogenous Myc expression may be acutely, systemically and reversibly hypomorphed to a degree broadly concordant with the ~30-60% reduced levels of Myc present in tissues of *Myc^+/−^* haploinsufficient mice ^57^ and mice in which key *Myc* enhancer elements have been deleted ^17^. We do see some modest variability across tissues in the extent of Myc hypomorphism, probably due to cell type variations in intrinsic activity of the *β-actin* “house-keeping” promoter we used to drive transgenic expression of the tTS^*Kid*^ repressor ^59^ and likely circumventable by driving expression from a ubiquitously expressed endogenous promoter ^60,61^.

Acute blunting of endogenous Myc expression in adult mice profoundly retarded development of KRas^*G12D*^-driven lung and pancreas tumours. Such protection is consistent with the reduced tumour incidence reported for germline *Myc* hemizygous and enhancer-deleted mice but even more dramatic.

Whereas activation of KRas^*G12D*^ in adult lung epithelium of p53-competent control animals typically drove sporadic development of multiple adenomas and adenocarcinomas with a spectrum of severity over our 32-week experimental timeframe, Myc-hypomorphed mice remained essentially tumour-free with lesions arrested at the stage of focal hyperplasia (AAH) for the entire duration save for a very few low-grade 1 lesions. Similar remarkable tumour suppression was also observed in p53-deficient KRas^*G12D*^-driven lung and pancreatic neoplasia. p53-deficient control (non-hypomorphed) mice developed widespread invasive lung or pancreatic tumours but the vast majority of focal lesions in the hypomorphed cohorts stalled as, respectively, focal AAH in lung or PanIN in pancreas. However, superimposed on this broad background of stalled pre-tumours we noted a small number of “escapee” lesions that had progressed and expanded into invasive adenocarcinomas in both lung and pancreas hypomorphed p53-deficient models. Virtually all of these rare “escapee” p53-deficient adenocarcinomas revealed underlying mechanisms that thwarted our switchable hypomorph model: either significant upregulation of Myc or loss of expression of the tTS^*Kid*^ repressor, or both - exceptions proving the rule that maximal, non-hypomorphed Myc levels are required for adenocarcinoma development. Hence, p53 inactivation does not defeat the powerful cancer protection afforded by Myc hypomorphism: rather, it exposes a vulnerability in the artificial mechanism we use to impose that hypomorphism. In this regard it is noteworthy that we never observed any escapee tumours in Myc hypomorphed p53-competent mice over the entire 32-week duration of our study, suggesting that the rate of sporadic inactivation of p53 in the hypomorphism-stalled lesions is negligible.

Our data indicate that Myc hypomorphism stalls cancer evolution at a critical bottleneck early in cancer evolution that governs the transition from indolent hyperplasia to overt neoplasia. To pinpoint the Myc dependency of this evolutionary bottleneck we asked the counter question - what level of Myc is required to drive stalled hyperplasias through the bottleneck? We recently demonstrated that deregulated Myc, expressed at only quasi-physiological levels, is sufficient to drive immediate and synchronous transition of stalled, indolent KRas^*G12D*^-driven lung AAHs or pancreatic PanINs to, respectively, aggressive and invasive lung and pancreatic adenocarcinomas. Remarkably, however, a modest 50% reduction in the level of deregulated Myc expression, broadly equivalent to the reduction in endogenous Myc expression induced by our switchable hypomorphism model and still sufficient to drive measurable transcriptional activity, was completely inert, driving neither transition from indolent pre-cancer to overt neoplasia nor detectable. Hence, both the capacity for endogenous Myc to support early tumour evolution and the capacity of deregulated Myc to force transition across the same evolutionary bottleneck exhibit the same tight dependency on a minimum threshold Myc expression and are profoundly blocked by Myc hypomorphism.

Acutely Myc hypomorphed adult animals showed no obvious short-term deleterious impact on normal adult physiology and health, begging the question of why evolution should have set endogenous Myc expression at levels that appear both “unnecessarily” high and pro-neoplastic to boot. To address this conundrum, we first used the temporal control afforded by our switchable hypomorph model to actively search for physiological processes that are sensitive to Myc hypomorphism. Myc is essential for successful transit through mid-gestation, a highly proliferative stage at which widespread co-expression of the Myc paralogue NMyc subsides ^62^ and subsequent tissue growth and maturation switches to sole dependence on Myc. *Myc*^−/−^ embryos die around 10-11 *dpc* due to widespread organogenic failure, most notably in vasculogenesis and haematopoiesis ^56^. Similarly, sustained imposition of Myc hypomorphism from conception in *MR* mice induced fully penetrant embryo failure, which occurred somewhat later than in *Myc^−/−^* embryos but shares its lethal hypovascular phenotype. Such ubiquitous mid-term lethality of actively hypomorphed *MR* embryos is not seen in germline haploinsufficient *Myc^+/−^* mice, almost all of which survive to adulthood. The reason for this difference is unclear but it underscores the distinction between passive germline *Myc* gene insufficiency and our active suppression of *Myc* expression, each of which likely perturbs in different ways the dynamics and kinetics of *Myc* regulation by mitogenic signals. Surprisingly, the failure specifically of hypomorphed *MR* embryos was observed only when the tTS^*Kid*^ repressor was contributed by the fathers: when mothers contributed the repressor, and were therefore themselves hypomorphed during pregnancy, no offspring of either *MR* or control genotypes were recoverable. Female infertility has also been reported in *Myc^+/−^* haploinsufficient mice, although with markedly less penetrance, and we guess indicates failure of some early Myc-dependent maternal function such as implantation, decidualization, or endometrial proliferation or angiogenesis ^58^. Whatever the proximal cause of infertility of hypomorphed *MR* mothers, however, reversal of hypomorphism rapidly returned all females to full fertility.

Although dependency on maximal Myc first arises during the intense proliferative burst in mid gestation, similarly high levels of proliferation continue in various organs throughout later gestation and into neonatal growth. Nonetheless, imposition of sustained Myc hypomorphism in either *MR* embryos post 13.5 *dpc* or in neonatal *MR* mice had no discernible impact on any aspect of development or on proliferation in any developing organ. Pups that were continuously hypomorphed from birth developed normally, at a rate indistinguishable from that of control littermates, achieving normal adult size and weight at the same time as their control littermates.

While the iatrogenic consequences of acute partial Myc inhibition appear surprisingly few and confined to mid-term gestation and female fertility, modulating Myc for cancer prophylaxis will most likely require systemic Myc inhibition over the long term, possibly years. Therefore, to model this we subjected adult *MR* mice to extended Myc hypomorphism. This uncovered a delayed but significant impact on haematopoiesis, starting with progressive onset leukopenia from around 6 weeks and consistent with the known obligate role of Myc in adult haematopoiesis, in particular in the maturation of haematopoietic stem cells ^63,64^. From around 12 weeks mice exhibit mild erythropenia, followed later by thrombocytosis (18 weeks), both consistent with the established role Myc plays in promoting erythropoiesis over megakaryocytopoiesis ^52^. These blood changes were accompanied by the development of modest splenic extramedullary haematopoiesis in around half the animals. However, all of these haematopoietic pathologies rapidly reversed upon relaxation of Myc hypomorphism. On this basis, we subjected *MR* mice to repeated metronomic Myc hypomorphism (4 weeks Myc suppression interspersed with 1-week recovery). This completely abrogated all the haematopoietic pathologies yet proved fully effective at blocking progression of KRas^*G12D*^-driven lung tumours.

The extent to which pharmacological partial Myc suppression in humans will replicate the cancer prophylaxis we see in mice awaits clarification. While no specific Myc inhibitor has yet been devised, valid experimental pharmacological strategies for indirectly inhibiting Myc already exist. All such current agents are relatively non-specific, targeting Myc transcription, translation and stability ^65^, and typically suppress Myc only transiently and partially and with significant side effects. As such, their utility in cancer therapy, where sustained complete Myc inhibition will likely be necessary, has been disappointing. Nonetheless, the partial and transient inhibition of Myc afforded by such agents might yet prove sufficient for cancer prophylaxis so long as a tolerable regimen could be found that balanced prophylactic efficacy with minimal side effects.

## SUPPLEMENTARY FIGURE LEGENDS

**Supplementary Figure 1:**
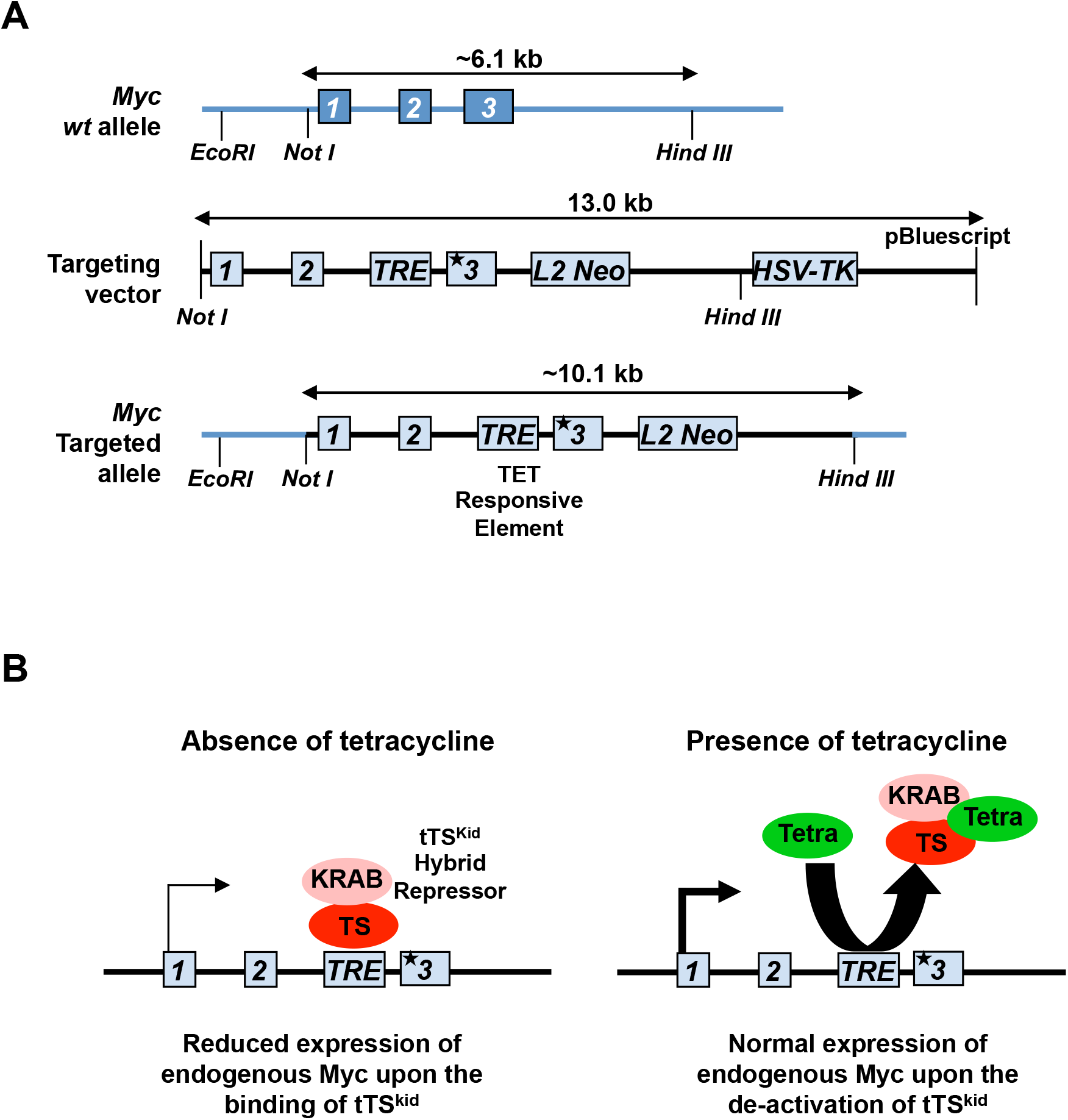
Design of the reversibly switchable Myc hypomorph allele. **A.** Schematic of wild type endogenous *Myc* gene, the targeting vector and the resultant *Myc* targeted allele (not to scale) with *TRE* insertion in intron 2. “*3” refers to silent mutations incorporated into exon 3 of the targeted *Myc* allele to distinguish it from that of the endogenous wild type Myc. **B.** Schematic of mechanism of action of reversible tTS^*Kid*^-regulated hypomorphism of the endogenous the *Myc* gene by tetracycline.

**Supplementary Figure 2:**
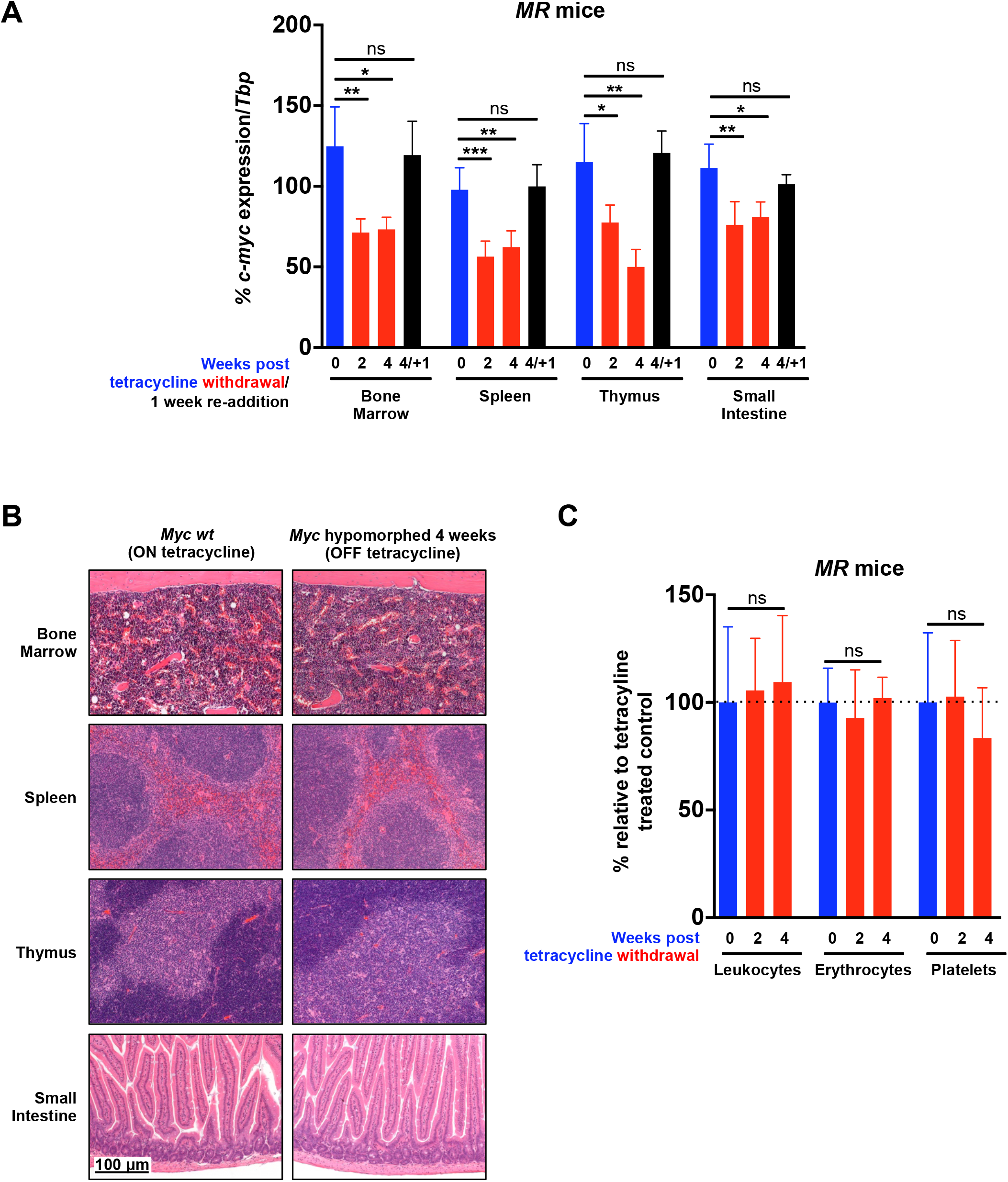
Hypomorphism of endogenous Myc imposed in adult mice has insignificant impact on proliferative tissues. **A.** Quantitative RT-PCR analysis of Myc mRNA expression in bone marrow, spleen, thymus, and small intestine from adult MR mice without tetracycline (i.e. Myc hypomorphed) for 0 (blue), 2, and 4 weeks (red), or without tetracycline for 4 weeks followed by restoration of tetracycline for 1 week (black). *Tbp* mRNA was used as a control gene. Results depict *Myc* expression mean + SD; n= 3-6 mice per treatment group. **B.** Representative H&E-stained sections of bone marrow, spleen, thymus and small intestine from control, tetracycline treated (non-hypomorphed) or 4-week Myc hypomorphed *MR* mice. **C.** Peripheral blood was collected from tetracycline treated (blue) or tetracycline deprived (red) MR mice. Data depict mean +SD of blood counts for leukocytes (left), erythrocytes (middle) and platelets (right) relative to tetracycline treated mice. n≥4 mice per treatment group. The unpaired t-test was used to analyze data. *p < 0.05, **p < 0.01, ***p < 0.001, ns = non-significant, SD= Standard deviation.

**Supplementary Figure 3:**
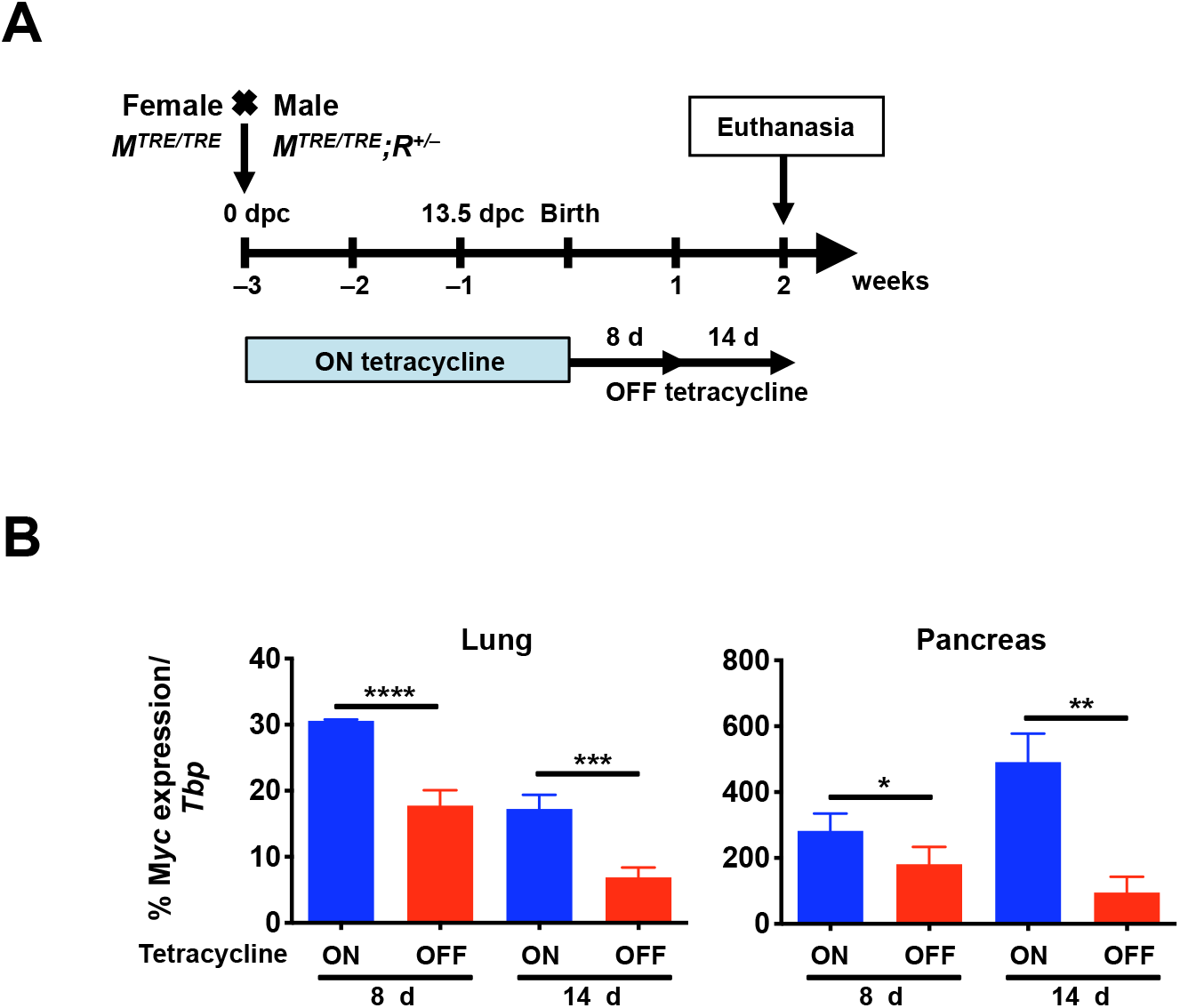
Induction of Myc hypomorphism in neonatal lung and pancreas. **A.** Schematic representation of study to determine impact of post-partum imposition of Myc hypomorphism on subsequent neonatal lung and pancreas development. *M^TRE/TRE^* females were mated with *M^TRE/TRE^;R^+/−^* males and pregnant females were maintained from conception on tetracycline until birth to maintain normal endogenous Myc levels. From birth (day 0) Myc was hypomorphed by cessation of tetracycline administration and neonatal mice were euthanized at 8 or 14 days of age. Control neonates maintained on tetracycline were euthanized at the same time points. **B.** Partial suppression of Myc expression in neonatal lung and pancreas. Quantitative RT-PCR analysis of *Myc* mRNA isolated from lungs (left) and pancreas (right) of 8 and 14 day-old control non-hypomorphed *M^TRE/TRE^;R^+/−^* (blue) versus *M^TRE/TRE^;R^+/−^* neonates hypomorphed from birth (red). Results depict mean + SD. *Tbp* was used as a control “housekeeping” gene. The unpaired t-test was used to analyze Taqman expression data. n =3-5 mice per treatment group. *p < 0.05, **p < 0.01, ***p < 0.001, ****p < 0.0001. SD = standard deviation.

**Supplementary Figure 4:**
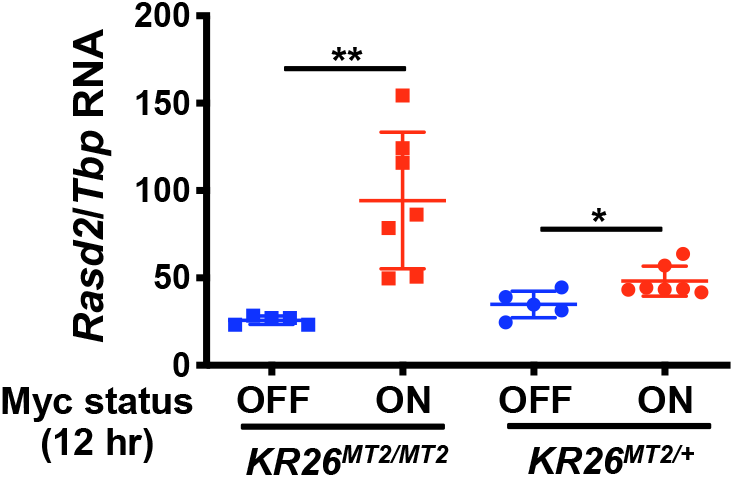
Hypomorphic Myc levels insufficient to drive transition to adenocarcinoma nonetheless retain measurable transcriptional activity. Quantitative RT-PCR was used to quantitate *Rasd2* mRNA expression in pancreata from 12-week old *KCR26^MT2/ MT2^* and *KCR26^MT2/+^* mice treated for 12 hrs with either oil control (MycER^T2^ OFF) or tamoxifen (MycER^T2^ ON). *Tbp* mRNA was used as a control gene. Results depict mean ± SD; n= 5-7 mice per treatment group. The unpaired student t-test was used to analyze Taqman expression data. *p < 0.05, **p < 0.01. SD= Standard deviation.

**Supplementary Figure 5:**
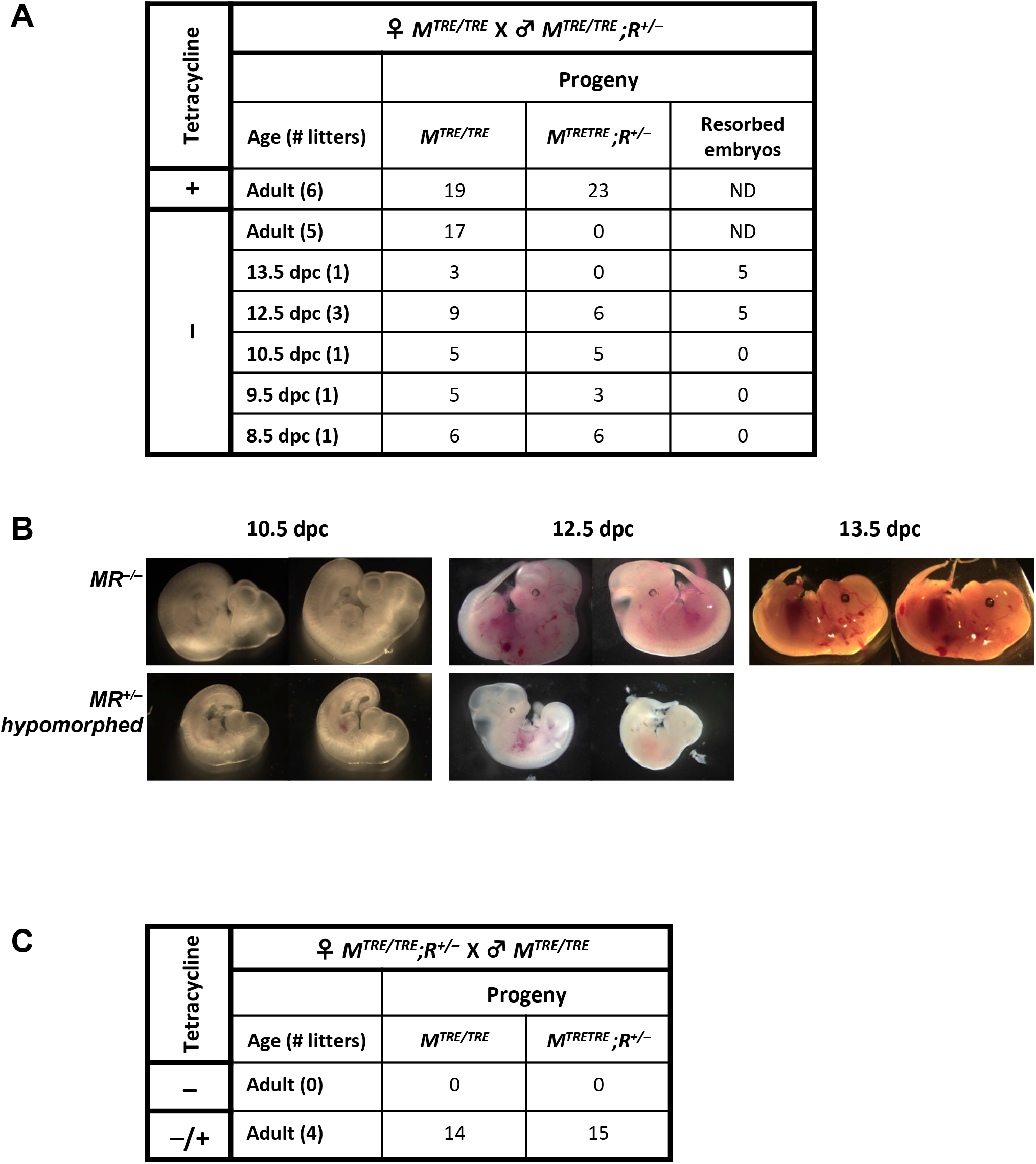
Imposition of Myc hypomorphism from conception triggers embryo failure. **A.** Table of number, fate and corresponding genotypes of embryos/pups derived by crossing *M* (*M^TRE^/M^TRE^*) females with *MR* (*M^TRE^/M^TRE^*;*R^+/−^*) males and collected at 8.5, 9.5, 10.5, 12.5, and 13.5 *dpc*, and post-weaning. **B.** Representative images of 10.5, 12.5 and 13.5 *dpc* littermate embryos derived by crossing *M* (*M^TRE^/M^TRE^*) females with *MR* (*M^TRE^/M^TRE^*;*R^+/−^*) males as described above (A) and maintained from conception in a Myc-hypomorphed state (the absence of tetracycline). **C.** Surviving embryos and corresponding genotypes of embryos/pups resulting from ♀*MR* (*M^TRE^/M^TRE^*;*R^+/−^*) X ♂*MR* (*M^TRE^/M^TRE^*) crosses in the absence of tetracycline (–) and after subsequent tetracycline re-administration (–/+). Hypomorphed MR Females were put OFF tetracycline 4 weeks before they were bred with M males. They were then maintained OFF tetracycline for another 6 weeks before they were put back ON tetracycline. MR females became pregnant about 4 weeks post-tetracycline re-administration.

**Supplementary Figure 6:**
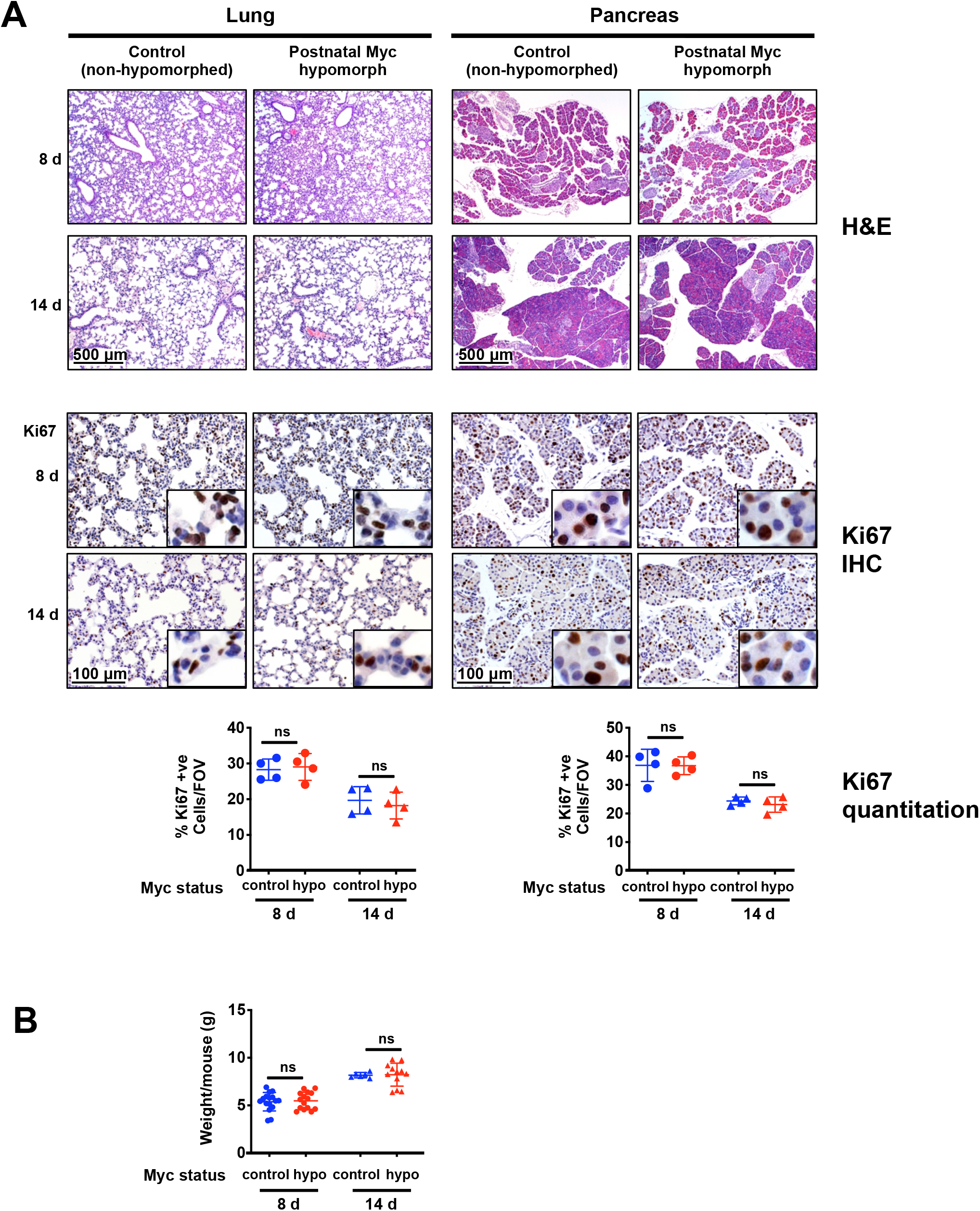
Post-partum imposition of Myc hypomorphism has no discernible impact on postnatal lung or pancreas architecture or proliferation, nor on overall neonatal growth. **A.** Top: Representative H&E-stained sections of lung and pancreas harvested from control versus hypomorphed neonates at 8 and 14 days post-partum. Middle: Representative Ki67 IHC-stained sections harvested from control versus hypomorphed neonates at 8 and 14 days post-partum, as described above. Bottom: Relevant quantification of IHC for Ki67 in sections of lung and pancreas harvested from control versus hypomorphed neonates at 8 and 14 days post-partum. Results represent mean ± SD, n = 4 mice per treatment group. **B.** Weights of 8 and 14 day old non-hypomorphed control *M^TRE/TR^;R^+/−^* (blue) versus *M^TRE/TRE^;R^+/−^* neonates hypomorphed from birth (red). Data depicts individual weights ± SD. The unpaired t-test was used to analyze data. ns = non-significant.

**Supplementary Figure 7:**
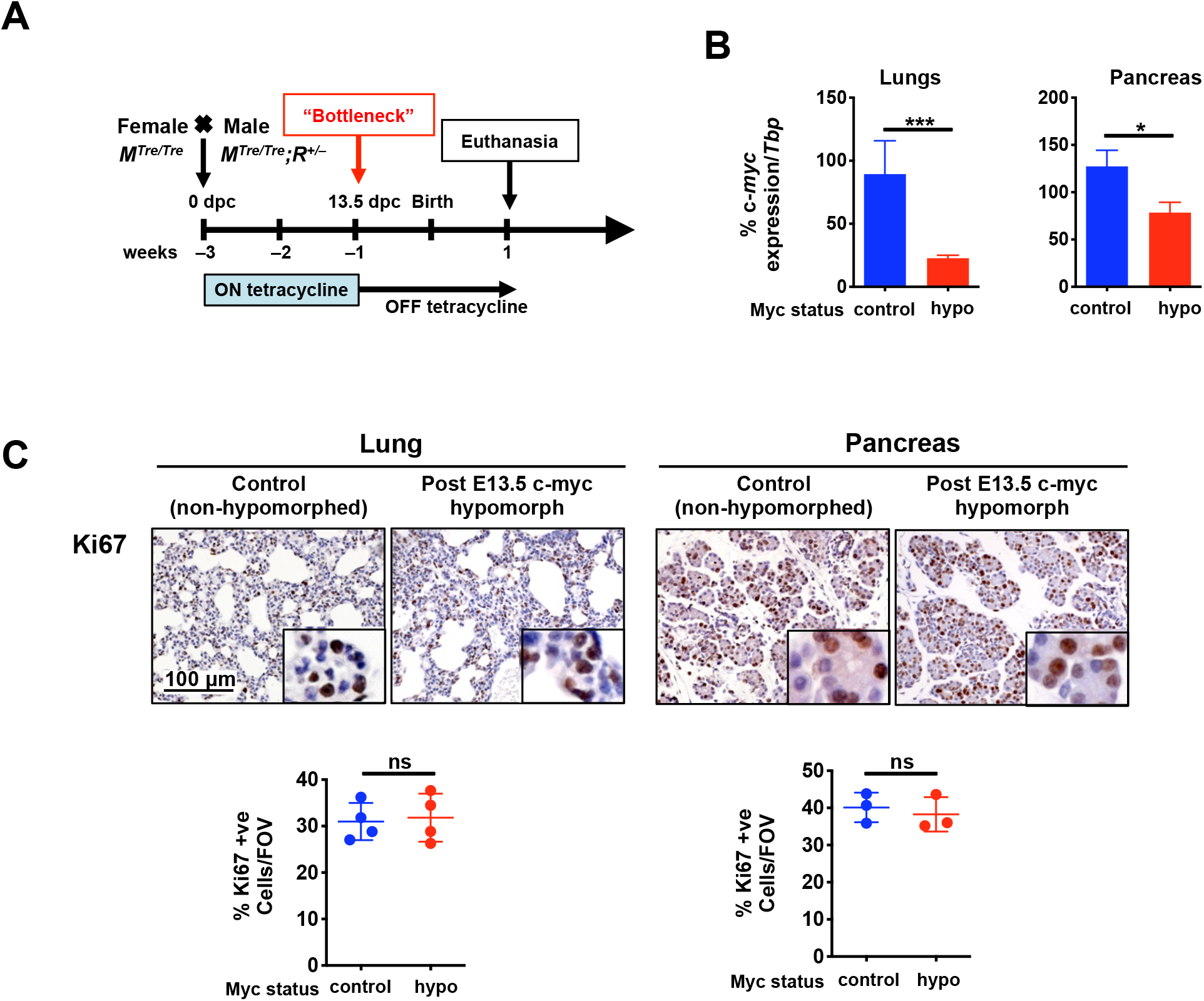
Imposition of Myc hypomorphism immediately after 13.5 *dpc* has no discernible impact on subsequent development or neonatal growth. **A.** Schematic of study. Embryos from *M^TRE/TRE^* females crossed with *M^TRE/TRE^;R^+/−^* males were maintained on tetracycline (normal Myc levels) until 13.5 *dpc*. Tetracycline was then withdrawn to induce Myc hypomorphism and neonates euthanized 8 days later. Control embryos were maintained on tetracycline throughout. **B.** Quantitative RT-PCR analysis of *Myc* mRNA isolated from lungs and pancreas of 8 day-old *M^TRE/TRE^;R^+/−^* neonates hypomorphed post 13.5 *dpc*, versus 8 day old control *M^TRE/TRE^;R^+/−^* mice maintained on tetracycline. Results depict mean + SD. *Tbp* was used as a control gene. n =3-5 mice per treatment group. **C.** Representative H&E stained and Ki67 IHC stained sections of lungs and pancreas from neonates described in (B). Below: Quantification of Ki67 IHC in neonatal lung and pancreas of mice hypomorphed post 13.5 *dpc*. Results represent mean ± SD, n = 3-4 mice per treatment group. The unpaired t-test was used to analyze data. *p < 0.05, ***p < 0.001, ns = non-significant. SD = standard deviation.

**Supplementary Figure 8:**
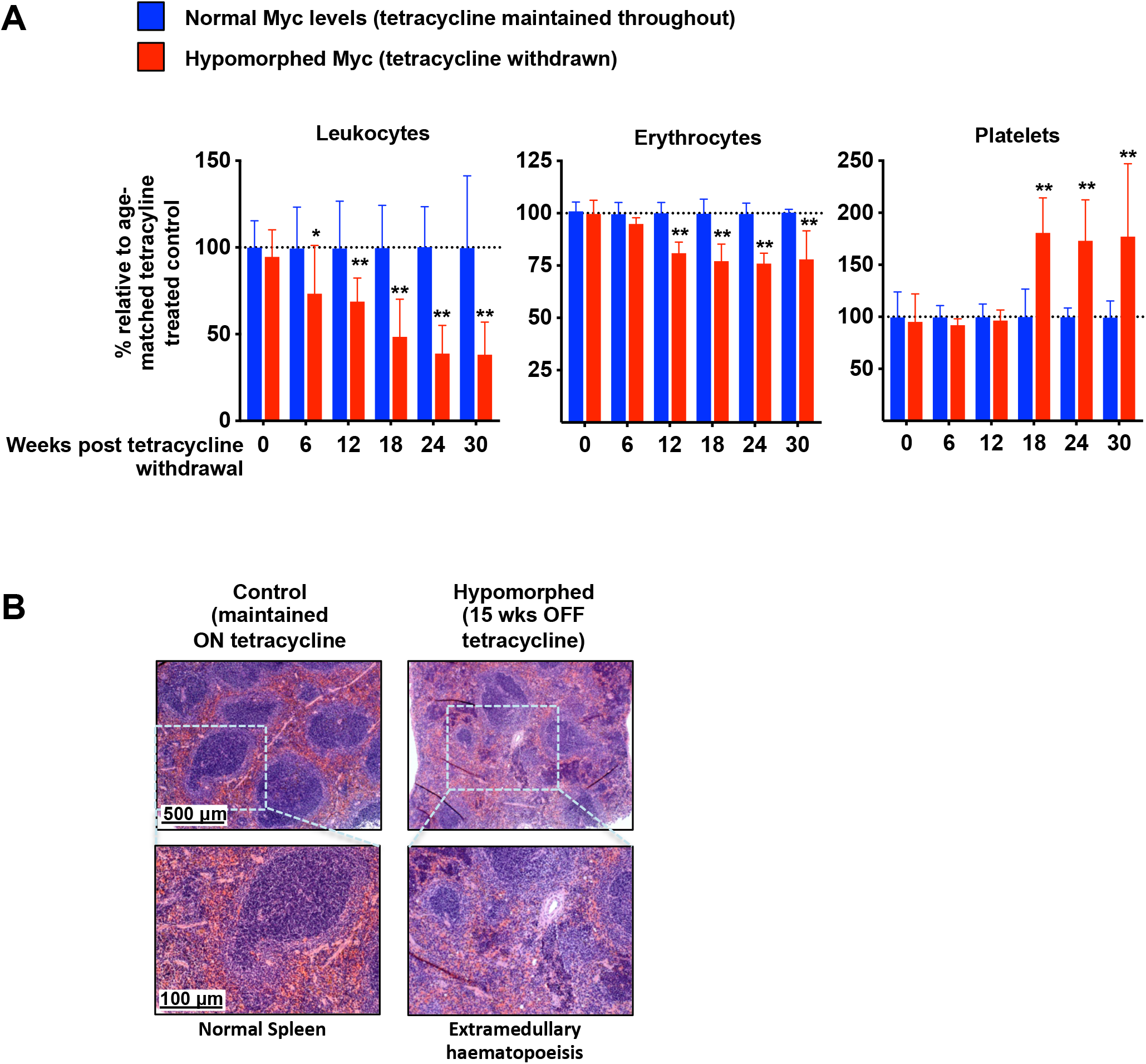
Full physiological Myc levels are required for long-term maintenance of haematopoiesis. **A.** *MR* mice were maintained on continuous tetracycline through development up to 5 weeks post-partum. Tetracycline was then withdrawn to hypomorph Myc and peripheral blood serially collected every 6 weeks thereafter (red). Control mice were maintained on tetracycline throughout (blue). The numbers of leukocytes (left), erythrocytes (middle), and platelet (right) are expressed relative to age-matched tetracycline treated controls. Results depict mean + SD. n≥3 mice per treatment group. *p < 0.05, **p < 0.01. SD = standard deviation. **B.** Representative H&E-stained sections of spleen from control *MR* mice (maintained on tetracycline) versus 15 weeks following tetracycline withdrawal.

## MATERIALS AND METHODS

### Generation and maintenance of genetically engineered mice

*p53^flox^; LSL-Kras^G12D^*, *Pdx-1-Cre*, *LSL-p53^R172H^, Rosa26-LSL-MER^T2^* and *β-actin-tTS* mice have all been previously described ^13,41,44,66–68^. To generate of *Myc^TRE^* mice, a targeting vector was constructed (Supplementary Figure 1) to include a heptameric tetracycline-response element (*TRE*) derived from the pTRE2 vector (Clontech), inserted into the intron 2 of the endogenous Myc locus that serves as a binding site for tetracycline-regulatable transcriptional repressors; silent mutations incorporated into exon3 (exon3 WT-GAAGAAGAG-; exon3 Mut-GA**G**GA**G**GA**A**-) to distinguish it from endogenous exon 3 of Myc, *LoxP*-flanked Neomycin resistance cassette L2 Neomycin used for positive selection; and a HSV-TK cassette added at the 3’end of the targeting vector to allow negative selection in mouse embryonic cells. Mouse embryonic stem cells (mESCs), from which *Myc^TRE^* mice were derived, were produced via homologous recombination of the endogenous Myc locus.

Mice were maintained on a 12-hour light/dark cycle with continuous access to food and water and in compliance with protocols approved by the UK Home Office guidelines under a project license to G.I.E. at the University of Cambridge. For Adenovirus-Cre recombinase (AdV-Cre) delivery, 8-10 week old mice were anaesthetized with isoflurane (Zoetis, IsoFlo 250 ml) and 5×10^7^ p.f.u. (plaque-forming units) of AdV-Cre were administered as described previously ^69^. Deregulated Myc activity was engaged in lung epithelia of *KM* mice or pancreatic epithelia of *KM^pdx1^* mice by daily intraperitoneal (i.p.) administration of tamoxifen (Sigma-Aldrich, TS648) dissolved in sunflower oil at a dose of 1 mg/20 g body mass; sunflower oil carrier was administered to control mice. *In vivo*, tamoxifen is metabolized by the liver into its active ligand 4-Hydroxytamoxifen (4-OHT). For long-term treatment, mice were placed on tamoxifen diet (Harlan Laboratories UK, TAM400 diet); control mice were maintained on regular diet. For Myc hypomorphism studies, *Myc^TRE/TRE^* mice were crossed into the *β-actin-tTS^Kid^* mice that ubiquitously express a chimeric transcriptional repressor molecule (tTS^*Kid*^) that can reversibly inhibit the expression of Myc endogenous gene. To keep systemic Myc at normal physiological level, *Myc^TRE/TRE^*; *β-actin-tTS^Kid/^*/– (*MR*) mice were maintained on drinking water containing tetracycline hydrochloride (100 mg/L) (Sigma-Aldrich, T7660) and 3% sucrose (Sigma-Aldrich, S9378) to increase palatability. To induce Myc hypomorphism, *MR* mice were transferred onto 3% sucrose drinking water.

### Embryo isolation

Embryos from timed MR matings (♀*M^TRE/TRE^* X ♂*M^TRE/TRE^;R^+/−^*) were isolated, dissected and rinsed with PBS at 4°C. Images were acquired using an M50 Leica Stereo microscope.

### Tissue preparation and histology

Tissues were isolated, fixed overnight in 10% neutral-buffered formalin, (Sigma-Aldrich, 501320), stored in 70% ethanol and paraffin embedded. Tissue sections (4 µM) were stained with hematoxylin and eosin (H&E) using standard reagents and protocols. Peripheral blood (50-80 µl) from the tail vein was collected in K_2_EDTA-coated BD Microtainer™ tubes (Fisher, 10346134) and analyzed at the Central Diagnostic Services, Queen’s Vet School Hospital, University of Cambridge.

### Immunohistochemistry and RNA scope

For immunohistochemistry (IHC), paraffin-embedded sections were de-paraffinized, rehydrated, and either boiled in 10 mM citrate buffer (pH 6.0) for Ki67 staining or in Tris/EDTA buffer (pH9.0) for Myc staining. Primary antibodies used were as follows: rabbit monoclonal anti-Ki67 (clone SP6) (Lab Vision, Fisher, 12603707) and rabbit monoclonal anti-Myc (Y69) (Abcam, ab32072). Primary antibodies were incubated with sections overnight at 4°C and detected using Vectastain Elite ABC HRP Kits (Vector Laboratories: Peroxidase Rabbit IgG PK-6101) and DAB substrate (Vector Laboratories, SK-4100); slides were then counterstained with hematoxylin solution (Sigma-Aldrich, GHS232). RNA-*in situ* hybridization (RNA-ISH) for *tTS^Kid^* was performed with a custom-designed probe, amplified with RNAscope 2.5 HD Reagent Kit (Advanced Cell Diagnostics; 322300) and developed with the TSA Plus kit (PerkinElmer; NEL760001KT) according to manufacturer’s instructions. Images were collected with a Zeiss Axio Imager M2 microscope equipped with Axiovision Rel 4.8 software.

### Quantitative real time-PCR

Total RNA was isolated from snap-frozen (liquid nitrogen) tissues using a Qiagen RNeasy Plus Isolation Kit followed by cDNA synthesis (High-Capacity cDNA RT kit, Applied Biosystems, 4374966). RT-PCR was performed using TaqMan Universal Master Mix II (Fisher, 4440038), according to manufacturer’s protocol. Primers used were: *Rasd2* (Fisher, Mm04209172_m1) and *Tbp* (Fisher, Mm00446973_m1). Samples were analyzed in triplicate on an Eppendorf Mastercycler Realpex 2 with accompanying software. *Tbp* was used as an internal amplification control.

### Quantification and Statistical Analysis

For quantification of tumour burden, H&E sections were scanned with an Aperio AT2 microscope (Leica Biosystems) at 20X magnification (resolution 0.5 microns per pixel) and analyzed with Aperio Software. IHC stained slides were quantified using Fiji open-source software. Statistical significance was assessed by unpaired t-test, with the mean values and the standard deviation (SD) calculated for each group using Prism GraphPad software. Unpaired t-test was also employed in RT-PCR statistical analysis. p values (ns = non-significant; *p < 0.05, **p < 0.01, ***p < 0.001, ****p < 0.0001) for each group.

## Author contributions

NMS and GIE conceived the project. GIE supervised the study with help from LBS and TDL. NMS designed experiments with help from LP. NMS and LP performed all experiments. RMK, YWK, SK, DG, AP and PA assisted with some experiments. NMS and GIE wrote the manuscript with help from LP. TDL edited the manuscript. All authors discussed results and revised the manuscript.

## Acknowledgments

We thank the members of the Evan laboratory for invaluable discussion and advice. We also thank Stephanie Whike, Michaela Griffin and Deborah Breiner for assistance with histology and genotyping. The study was supported by programme grants (Cancer Research UK C4750/A12077, C4750/A19013A and C4750/A29210, European Research Council Advanced Investigator Award (294851), and a Stand Up To Cancer-Cancer Research UK-Lustgarten Foundation Pancreatic Cancer Dream Team Research Grant (Grant Number: SU2C-AACR-DT20-16), all to GIE. Stand Up To Cancer is a division of the Entertainment Industry Foundation and the research grant is administered by the American Association for Cancer Research, the Scientific Partner of SU2C. GIE is a member of AstraZeneca’s IMED oncology external science advisory panel.

## References

1 Sodir, N. M. et al. Endogenous Myc maintains the tumor microenvironment. Genes & Development 25, 907–916, doi:papers3://publication/doi/10.1101/gad.2038411 (2011).

2 Soucek, L. et al. Modelling Myc inhibition as a cancer therapy. Nature 455, 679–683 (2008).

3 Whitfield, J. R., Beaulieu, M. E. & Soucek, L. Strategies to Inhibit Myc and Their Clinical Applicability. Front Cell Dev Biol 5, 10, doi:10.3389/fcell.2017.00010 (2017).

4 Sodir, N. M. & Evan, G. I. Finding cancer’s weakest link. Oncotarget; Vol 2, No 12: December 2011 (2011).

5 Facchini, L. M., Chen, S., Marhin, W. W., Lear, J. N. & Penn, L. Z. The Myc negative autoregulation mechanism requires Myc-Max association and involves the c-*myc* P2 minimal promoter. Molecular and Cellular Biology 17, 100–114, doi:papers3://publication/doi/10.1128/mcb.17.1.100 (1997).

6 Grignani, F. et al. Negative autoregulation of c-*myc* gene expression is inactivated in transformed cells. EMBO J 9, 3913–3922, doi:papers3://publication/uuid/7039DA13-607B-4D84-B054-A3F57446DBE0 (1990).

7 Penn, L. J., Brooks, M. W., Laufer, E. M. & Land, H. Negative autoregulation of c-*myc* transcription. EMBO J 9, 1113–1121, doi:papers3://publication/uuid/2828050D-6B98-4C25-8800-B42FA02C904A (1990).

8 Dean, M. et al. Regulation of c-*myc* transcription and mRNA abundance by serum growth factors and cell contact. J. Biol. Chem. 261, 9161–9166 (1986).

9 Waters, C. M., Littlewood, T. D., Hancock, D. C., Moore, J. P. & Evan, G. I. c-myc protein expression in untransformed fibroblasts. Oncogene 6, 797–805 (1991).

10 Dang, C. V. MYC on the Path to Cancer. Cell 149, 22–35, doi:papers3://publication/doi/10.1016/j.cell.2012.03.003 (2012).

11 Lin, C. Y. et al. Transcriptional amplification in tumor cells with elevated c-Myc. Cell 151, 56–67, doi:10.1016/j.cell.2012.08.026 (2012).

12 Lossos, I. S. et al. Transformation of follicular lymphoma to diffuse large-cell lymphoma: Alternative patterns with increased or decreased expression of c-*myc* and its regulated genes. Proceedings of the National Academy of Sciences 99, 8886, doi:10.1073/pnas.132253599 (2002).

13 Murphy, D. J. et al. Distinct Thresholds Govern Myc’s Biological Output In Vivo. Cancer Cell 14, 447–457, doi:papers3://publication/doi/10.1016/j.ccr.2008.10.018 (2008).

14 Kortlever, R. M. et al. Myc Cooperates with Ras by Programming Inflammation and Immune Suppression. Cell 171, 1301–1315.e1314, doi:papers3://publication/doi/10.1016/j.cell.2017.11.013 (2017).

15 Sodir, N. M. et al. MYC Instructs and Maintains Pancreatic Adenocarcinoma Phenotype. Cancer Discovery 10, 588–607, doi:papers3://publication/doi/10.1158/2159-8290.CD-19-0435 (2020).

16 Ahmadiyeh, N. et al. 8q24 prostate, breast, and colon cancer risk loci show tissue-specific long-range interaction with MYC. Proceedings of the National Academy of Sciences 107, 9742–9746, doi:papers3://publication/doi/10.1073/pnas.0910668107 (2010).

17 Dave, K. et al. Mice deficient of Myc super-enhancer region reveal differential control mechanism between normal and pathological growth. eLife 6, 9742, doi:papers3://publication/doi/10.7554/eLife.23382 (2017).

18 Al Olama, A. A. et al. Multiple loci on 8q24 associated with prostate cancer susceptibility. Nature Publishing Group, 1–3, doi:papers3://publication/doi/10.1038/ng.452 (2009).

19 Amundadottir, L. T. et al. A common variant associated with prostate cancer in European and African populations. Nature Publishing Group 38, 652–658, doi:papers3://publication/doi/10.1038/ng1808 (2006).

20 Gudmundsson, J. et al. Genome-wide association study identifies a second prostate cancer susceptibility variant at 8q24. Nature Publishing Group 39, 631–637, doi:papers3://publication/doi/10.1038/ng1999 (2007).

21 Yeager, M. et al. Genome-wide association study of prostate cancer identifies a second risk locus at 8q24. Nature Publishing Group 39, 645–649, doi:papers3://publication/doi/10.1038/ng2022 (2007).

22 CORGI, C. et al. A genome-wide association scan of tag SNPs identifies a susceptibility variant for colorectal cancer at 8q24.21. Nature Publishing Group 39, 984–988, doi:papers3://publication/doi/10.1038/ng2085 (2007).

23 Jia, Y., Chng, W.-J. & Zhou, J. Super-enhancers: critical roles and therapeutic targets in hematologic malignancies. Journal of Hematology & Oncology 12, 77, doi:10.1186/s13045-019-0757-y (2019).

24 Schuijers, J. et al. Transcriptional Dysregulation of MYC Reveals Common Enhancer-Docking Mechanism. Cell Rep 23, 349–360, doi:10.1016/j.celrep.2018.03.056 (2018).

25 Herranz, D. et al. A NOTCH1-driven MYC enhancer promotes T cell development, transformation and acute lymphoblastic leukemia. Nature Medicine 20, 1130–1137, doi:papers3://publication/doi/10.1038/nm.3665 (2014).

26 Zhang, X. et al. Identification of focally amplified lineage-specific super-enhancers in human epithelial cancers. Nature Genetics 48, 176–182, doi:10.1038/ng.3470 (2016).

27 Sur, I. K. et al. Mice Lacking a *Myc* Enhancer That Includes Human SNP rs6983267 Are Resistant to Intestinal Tumors. Science 338, 1360–1363, doi:10.1126/science.1228606 (2012).

28 Homer-Bouthiette, C. et al. Deletion of the murine ortholog of the 8q24 gene desert has anti-cancer effects in transgenic mammary cancer models. 1–16, doi:papers3://publication/doi/10.1186/s12885-018-5109-8 (2018).

29 Davis, A. C., Wims, M., Spotts, G. D., Hann, S. R. & Bradley, A. A null c-*myc* mutation causes lethality before 10.5 days of gestation in homozygotes and reduced fertility in heterozygous female mice. Genes & Development 7, 671–682, doi:papers3://publication/doi/10.1101/gad.7.4.671 (1993).

30 Trumpp, A. et al. c-Myc regulates mammalian body size by controlling cell number but not cell size. Nature 414, 768–773. (2001).

31 Yekkala, K. & Baudino, T. A. Inhibition of intestinal polyposis with reduced angiogenesis in ApcMin/+ mice due to decreases in c-Myc expression. Mol. Cancer Res. 5, 1296–1303, doi:papers3://publication/doi/10.1158/1541-7786.MCR-07-0232 (2007).

32 Athineos, D. & Sansom, O. J. Myc heterozygosity attenuates the phenotypes of APC deficiency in the small intestine. Oncogene 29, 2585–2590, doi:papers3://publication/doi/10.1038/onc.2010.5 (2010).

33 Walz, S. et al. Activation and repression by oncogenic MYC shape tumour-specific gene expression profiles. Nature, 1–18, doi:papers3://publication/doi/10.1038/nature13473 (2014).

34 Bazarov, A. V. et al. A modest reduction in c-myc expression has minimal effects on cell growth and apoptosis but dramatically reduces susceptibility to Ras and Raf transformation. Cancer Research 61, 1178–1186, doi:papers3://publication/uuid/949B9AEE-D1B9-43F4-B938-7321D3A3A0E0 (2001).

35 Mateyak, M. K., Obaya, A. J., Adachi, S. & Sedivy, J. M. Phenotypes of c-Myc-deficient rat fibroblasts isolated by targeted homologous recombination. Cell Growth Differ 8, 1039–1048. (1997).

36 Guney, I., Wu, S. & Sedivy, J. M. Reduced c-Myc signaling triggers telomere-independent senescence by regulating Bmi-1 and p16(INK4a). Science 103, 3645–3650, doi:papers3://publication/doi/10.1073/pnas.0600069103 (2006).

37 Clarke, A. R. Manipulating the germline: its impact on the study of carcinogenesis. Carcinogenesis 21, 435–441, doi:10.1093/carcin/21.3.435 (2000).

38 El-Brolosy, M. A. & Stainier, D. Y. R. Genetic compensation: A phenomenon in search of mechanisms. PLoS Genet 13, e1006780–1006717, doi:papers3://publication/doi/10.1371/journal.pgen.1006780 (2017).

39 Sage, J. et al. Targeted disruption of the three Rb-related genes leads to loss of G(1) control and immortalization. Genes & Development 14, 3037–3050, doi:papers3://publication/doi/10.1101/gad.843200 (2000).

40 Sheng, Y. et al. Role of c-Myc haploinsufficiency in the maintenance of HSCs. Blood, doi:10.1182/blood.2019004688 (2020).

41 Gamper, I. et al. Determination of the physiological and pathological roles of E2F3 in adult tissues. Sci. Rep., 1–15, doi:papers3://publication/doi/10.1038/s41598-017-09494-6 (2017).

42 Tekki-Kessaris, N., Bonventre, J. V. & Boulter, C. A. Characterization of the mouse *Kid1* gene and identification of a highly related gene, Kid2. Gene 240, 13–22 (1999).

43 Yaghmai, R. & Cutting, G. R. Optimized regulation of gene expression using artificial transcription factors. Mol Ther 5, 685–694 (2002).

44 Jackson, E. L. et al. Analysis of lung tumor initiation and progression using conditional expression of oncogenic K-*ras*. Genes Dev 15, 3243–3248 (2001).

45 Jackson, E. L. et al. The differential effects of mutant p53 alleles on advanced murine lung cancer. Cancer Res 65, 10280–10288 (2005).

46 Hingorani, S. R. et al. Trp53^*R172H*^ and Kras^*G12D*^ cooperate to promote chromosomal instability and widely metastatic pancreatic ductal adenocarcinoma in mice. Cancer Cell 7, 469–483 (2005).

47 Volckaert, T. & De Langhe, S. Lung epithelial stem cells and their niches: Fgf10 takes center stage. 7, 1–15, doi:papers3://publication/doi/10.1186/1755-1536-7-8 (2014).

48 Bonal, C. et al. Pancreatic inactivation of c-Myc decreases acinar mass and transdifferentiates acinar cells into adipocytes in mice. Gastroenterology 136, 309–319.e309, doi:papers3://publication/doi/10.1053/j.gastro.2008.10.015 (2009).

49 Nakhai, H., Siveke, J. T., Mendoza-Torres, L. & Schmid, R. M. Conditional inactivation of Myc impairs development of the exocrine pancreas. Development 135, 3191–3196, doi:papers3://publication/doi/10.1242/dev.017137 (2008).

50 Stellas, D. et al. Therapeutic effects of an anti-Myc drug on mouse pancreatic cancer. JNCI Journal of the National Cancer Institute 106, dju320–dju320, doi:papers3://publication/doi/10.1093/jnci/dju320 (2014).

51 Zhou, Q. et al. A multipotent progenitor domain guides pancreatic organogenesis. Developmental Cell 13, 103–114, doi:papers3://publication/doi/10.1016/j.devcel.2007.06.001 (2007).

52 Guo, Y. et al. c-Myc-mediated control of cell fate in megakaryocyte-erythrocyte progenitors. Blood 114, 2097–2106, doi:papers3://publication/doi/10.1182/blood-2009-01-197947 (2009).

53 Delgado, M. D. & Leon, J. Myc roles in hematopoiesis and leukemia. Genes & Cancer 1, 605–616, doi:papers3://publication/doi/10.1177/1947601910377495 (2010).

54 Baena, E., Ortiz, M., Martinez-A, C. & Moreno de Alborán, I. c-Myc is essential for hematopoietic stem cell differentiation and regulates Lin−Sca-1+c-Kit− cell generation through p21. Experimental Hematology 35, 1333–1343, doi:papers3://publication/doi/10.1016/j.exphem.2007.05.015 (2007).

55 de Alboran, I. M. et al. Analysis of C-MYC function in normal cells via conditional gene-targeted mutation. Immunity 14, 45–55, doi:papers3://publication/uuid/737A364C-BB8D-440A-83E6-4A659064BA86 (2001).

56 Baudino, T. A. et al. c-Myc is essential for vasculogenesis and angiogenesis during development and tumor progression. Genes & Development 16, 2530–2543, doi:papers3://publication/doi/10.1101/gad.1024602 (2002).

57 Hofmann, J. W. et al. Reduced Expression of MYC Increases Longevity and Enhances Healthspan. Cell 160, 477–488, doi:papers3://publication/doi/10.1016/j.cell.2014.12.016 (2015).

58 Ramathal, C., Bagchi, I., Taylor, R. & Bagchi, M. Endometrial Decidualization: Of Mice and Men. Semin Reprod Med 28, 017–026, doi:papers3://publication/doi/10.1055/s-0029-1242989 (2010).

59 Ruan, W. & Lai, M. Actin, a reliable marker of internal control? Clin Chim Acta 385, 1–5, doi:10.1016/j.cca.2007.07.003 (2007).

60 Jacox, E., Gotea, V., Ovcharenko, I. & Elnitski, L. Tissue-specific and ubiquitous expression patterns from alternative promoters of human genes. PLoS ONE 5, e12274–e12274, doi:10.1371/journal.pone.0012274 (2010).

61 Lee, H. Genetically engineered mouse models for drug development and preclinical trials. Biomol Ther (Seoul) 22, 267–274, doi:10.4062/biomolther.2014.074 (2014).

62 Mugrauer, G., Alt, F. W. & Ekblom, P. N-myc proto-oncogene expression during organogenesis in the developing mouse as revealed by *in situ* hybridization. J Cell Biol 107, 1325–1335, doi:10.1083/jcb.107.4.1325 (1988).

63 Wilson, A. et al. c-Myc controls the balance between hematopoietic stem cell self-renewal and differentiation. Genes Dev 18, 2747–2763 (2004).

64 Delgado, M. D. & León, J. Myc roles in hematopoiesis and leukemia. Genes & Cancer 1, 605–616, doi:10.1177/1947601910377495 (2010).

65 Carabet, L., Rennie, P. & Cherkasov, A. Therapeutic Inhibition of Myc in Cancer. Structural Bases and Computer-Aided Drug Discovery Approaches. IJMS 20, 120–149, doi:papers3://publication/doi/10.3390/ijms20010120 (2019).

66 Hingorani, S. R. et al. Preinvasive and invasive ductal pancreatic cancer and its early detection in the mouse. Cancer Cell 4, 437–450 (2003).

67 Marino, S., Vooijs, M., van Der Gulden, H., Jonkers, J. & Berns, A. Induction of medulloblastomas in p53-null mutant mice by somatic inactivation of Rb in the external granular layer cells of the cerebellum. Genes Dev 14, 994–1004 (2000).

68 Olive, K. P. et al. Mutant p53 gain of function in two mouse models of Li-Fraumeni syndrome. Cell 119, 847–860, doi:10.1016/j.cell.2004.11.004 (2004).

69 Fasbender, A. et al. Incorporation of adenovirus in calcium phosphate precipitates enhances gene transfer to airway epithelia *in vitro* and *in vivo*. J Clin Invest 102, 184–193, doi:10.1172/jci2732 (1998).

